# Linkage-aware inference of fitness from short-read time-series genomic data

**DOI:** 10.64898/2026.01.08.698347

**Authors:** Syed Muhammad Umer Abdullah, Muhammad Saqib Sohail, Raymond H. Y. Louie, Yanni Sun, John P. Barton, Matthew R. McKay

## Abstract

Inferring the fitness effect of mutations is a basic problem in understanding the evolution of populations over time. When multiple mutations are present in a population simultaneously, genetic linkage comes into play, and the fate of an individual mutation depends on both its fitness as well as the background on which it occurs. Accurate inference of fitness effects for evolutionary systems with multiple competing mutations is therefore contingent on resolving the confounding effects of genetic linkage, captured by the covariance between allele-pairs. Increasingly, evolutionary studies are using short-read sequencing technologies to produce detailed snapshots of evolving populations. This presents a problem as the frequencies of allele-pairs are not known beyond the read-length, hampering any attempt to resolve the effects of genetic linkage between pairs of loci residing on different reads. Here we present a computationally efficient pipeline for inferring selection from short-read time-series data with partial allele-pair frequency information, while accounting for linkage. Simulation results show that the method has good performance and is scalable to systems with several thousand variants. Additionally, we demonstrate the pipeline’s utility on real datasets of within-host HIV and SARS-CoV-2 evolution, showcasing its applicability in resolving linkage effects from complex evolutionary histories.

## 1 Introduction

The evolution of a population (organisms of the same species living and reproducing in a particular environment) is shaped by the complex interplay of various evolutionary phenomenon such as selection, mutation, drift, recombination, genetic linkage, and others. Genetic linkage—the tendency of mutations to be inherited together—becomes an important factor in shaping evolution when multiple loci are simultaneously polymorphic in the population. Experimental and observational evidence shows that genetic linkage is an important evolutionary factor in several real evolving populations (Hegreness *et al*., 2006; Strelkowa and Lässig, 2012; Lang *et al*., 2013). Genetic linkage has been shown to affect drug resistance in the protozoan parasite Plasmodium falciparum (Alam *et al*., 2011), malaria protection in humans (Sanchez-Mazas *et al*., 2017), cancer progression (Lee *et al*., 2024), and evolution of viruses such as the human immunodeficiency virus (HIV) (Sohail *et al*., 2021) and influenza (Strelkowa & Lässig, 2012, Łuksza & Lässig, 2014). Genetic linkage may result in hitchhiking (Smith and Haigh, 1974), positive or negative (clonal) interference (Gerrish and Lenski, 1998) or background selection (Charlesworth *et al*., 1993; Charlesworth, 1994).

Researchers have long studied population-level time-series genomic data to identify and understand the evolutionary phenomenon responsible for the observed patterns of genetic variation. These efforts have been boosted by the developments in short-read sequencing technologies (Metzker, 2010; Elshire *et al*., 2011; Ozsolak and Milos, 2011; van Dijk *et al*., 2014; Goodwin *et al*., 2016) which have enabled the collection of high-resolution time-series genomic data. Examples of such data include evolve-and-resequence studies under lab settings of viruses (Cordey *et al*., 2010; Bertels *et al*., 2019), bacteria (Van Den Bergh *et al*., 2016; Barroso-Batista *et al*., 2020), yeast (Lang *et al*., 2013; Frenkel *et al*., 2014; Raynes *et al*., 2018; Fisher *et al*., 2018), and fruit flies (Orozco-Terwengel *et al*., 2012; Griffin *et al*., 2017), and studies of in vivo evolution of pathogens including HIV (Zanini *et al*., 2015; Setliff *et al*., 2018), influenza (Imai *et al*., 2012; McCrone *et al*., 2018; Xue *et al*., 2018), hepatitis C virus (Bull *et al*., 2011; Takeda *et al*., 2017), severe acute respiratory syndrome coronavirus 2 (SARS-CoV-2) (Boshier *et al*., 2022; Kemp *et al*., 2021) and cancer (Fischer *et al*., 2014; Ishaque *et al*., 2018). These and other similar evolutionary data provide an opportunity to observe evolving populations in remarkable detail.

Several methods exist in the literature that can use the mutant allele frequency trajectories observable in data obtained from short-read sequencing platforms to infer evolutionary parameters in scenarios where genetic linkage between loci is negligible (Bollback *et al*., 2008; Malaspinas *et al*., 2012; Mathieson and McVean, 2013; Feder *et al*., 2014; Lacerda and Seoighe, 2014; Foll *et al*., 2015; Topa *et al*., 2015; Ferrer-Admetlla *et al*., 2016; Schraiber *et al*., 2016; Iranmehr *et al*., 2017; Taus *et al*., 2017; Zinger *et al*., 2019). However, in populations where genetic linkage influences evolution, relying solely on mutant allele frequencies without considering interactions between alleles can lead to inaccurate inference of evolutionary parameters. Linkage-aware methods (He *et al*., 2020; Buffalo and Coop, 2019; Terhorst *et al*., 2015; Illingworth, 2015; Sohail *et al*., 2021; Shen *et al*., 2021) can analyze the evolution of multiple loci collectively to disentangle the effects of genetic linkage. However, despite their utility for analyzing linked loci (Terhorst *et al*., 2015; Illingworth 2015; Buffalo & Coop, 2019; He *et al*., 2020; Shen *et al*., 2021) these methods are limited to small system size (Terhorst *et al*., 2015; He *et al*., 2020), lack the ability to quantify fitness effect of individual mutations (Buffalo & Coop, 2019), or incur a high computational cost (Terhorst *et al*., 2015; Illingworth 2015; Buffalo and Coop, 2019; He *et al*., 2020; Shen *et al*., 2021). In contrast, Sohail *et al*., 2021 proposed marginal path likelihood (MPL), a fast and scalable framework that leads to closed-form analytical estimates of selection coefficients from time-series data while accounting for the effects of genetic linkage in addition to those of mutation, drift and recombination. MPL has found applications in different scenarios, such as fitness inference in the presence of sampling noise (Cheng *et al*., 2025), fitness inference in the presence of epistasis (Sohail *et al*., 2022), understanding HIV-1 escape (Gao and Barton, 2025), and analyzing SARS-CoV-2 data (Zeng H *et al*., 2024; Lee *et al*., 2025).

In the MPL framework, the linkage between multiple alleles is taken into account through the covariance matrix of the mutant allele frequencies at each time-point. The entries of this covariance matrix are a function of single and double mutant allele frequencies and are equivalent to a classical linkage disequilibrium metric (Hedrick, 1987). Inferring this covariance matrix is a significant challenge for short-read sequencing technologies, which are commonly used to observe evolving populations. While short-read sequencing provides accurate estimates of the single mutant allele frequencies, only the double mutant allele frequencies between allele pairs residing on the same read are directly observable. A recent approach (Li and Barton, 2023) presented a method to estimate mutant allele covariance matrices from the product of the change in single mutant allele frequencies. This approach is sensitive to the frequency of temporal sampling since changes in the single mutant allele frequencies become difficult to observe as the temporal sampling step increases. The methodological challenge of robustly inferring double mutant allele frequencies from shortread data across general evolutionary scenarios remains to be addressed. This work builds upon the MPL framework and presents a novel pipeline for the inference of selection from short-read time-series data. The proposed computational pipeline, termed maximum path likelihood with population reconstruction (MPL-R), takes in short-read time-series data, infers the double mutant allele frequencies for all allele-pairs beyond the read length (inter-read allele-pairs), and uses these to infer selection. The main idea is to reconstruct full-length sequences of an evolving population from short reads, from which the double mutant allele frequencies can be computed. However, population reconstruction is a challenging problem to solve (Posada-Cespedes *et al*., 2017). The reconstruction process may result in the reconstruction of erroneous haplotypes, especially those inferred to have a low frequency in the population and is particularly error-prone when the actual population consists of multiple haplotypes (Schirmer *et al*., 2014). We show in this work that for the purpose of inferring selection, it is not necessary to reconstruct the actual population exactly, and one only needs to obtain a probable set of sequences approximating the actual population. We use an off the shelf population reconstruction algorithm, Quasirecomb (Töpfer *et al*., 2013), and employ a novel reconstruction strategy based on bootstrap aggregating (bagging) (Breiman, 1996), which allows for leveraging parallel processing on sub-sampled data leading to reduced memory requirements and wall clock time without significantly affecting the accuracy of the final estimates. We demonstrate that Quasirecomb, while making some errors in reconstructing full populations, does an excellent job at producing accurate double mutant allele frequencies. We test the performance of MPL-R on simulated data and show that the pipeline accurately infers selection over a range of parameters. Interestingly, we found that inference using partial covariance information (e.g., using only covariance values that could be measured directly from short-read data) yields worse results than ignoring covariances entirely. Comparison with a simpler MPL implementation that naively employs observed double mutant allele frequencies while disregarding all unobservable double mutant allele frequencies (corresponding to allele-pairs residing on different reads), has vastly suboptimal performance. This emphasizes the importance of constructing double mutant allele frequencies both within and outside sequencing reads, as achieved with MPL-R. We further demonstrate the performance of MPL-R by applying it to genetic data from a cohort of HIV-1 infected patients, demonstrating that it can capture similar biological insights from short-read data as obtainable with full length sequences.

## 2 Methods

### 2.1 Overview

We present a pipeline that infers selection from short-read time-series data based on the MPL framework. The main challenge addressed in this work is the inference of inter-read covariance information (which is not directly observable from short-read data) required by the MPL framework. To infer the unobserved inter-read covariance information, MPL-R performs population reconstruction from short-read data at each sampled time-point using Quasirecomb (Töpfer *et al*., 2013). MPL-R employs a novel reconstruction strategy based on bootstrap aggregating (bagging) approach (Breiman, 1996). The single and double mutant allele frequencies are then computed from the reconstructed sequences, which are then used to obtain the estimates of selection coefficients adjusted for linkage effects. The workflow of the proposed MPL-R pipeline is shown in **Fig. 1**.

**Fig. 1.**
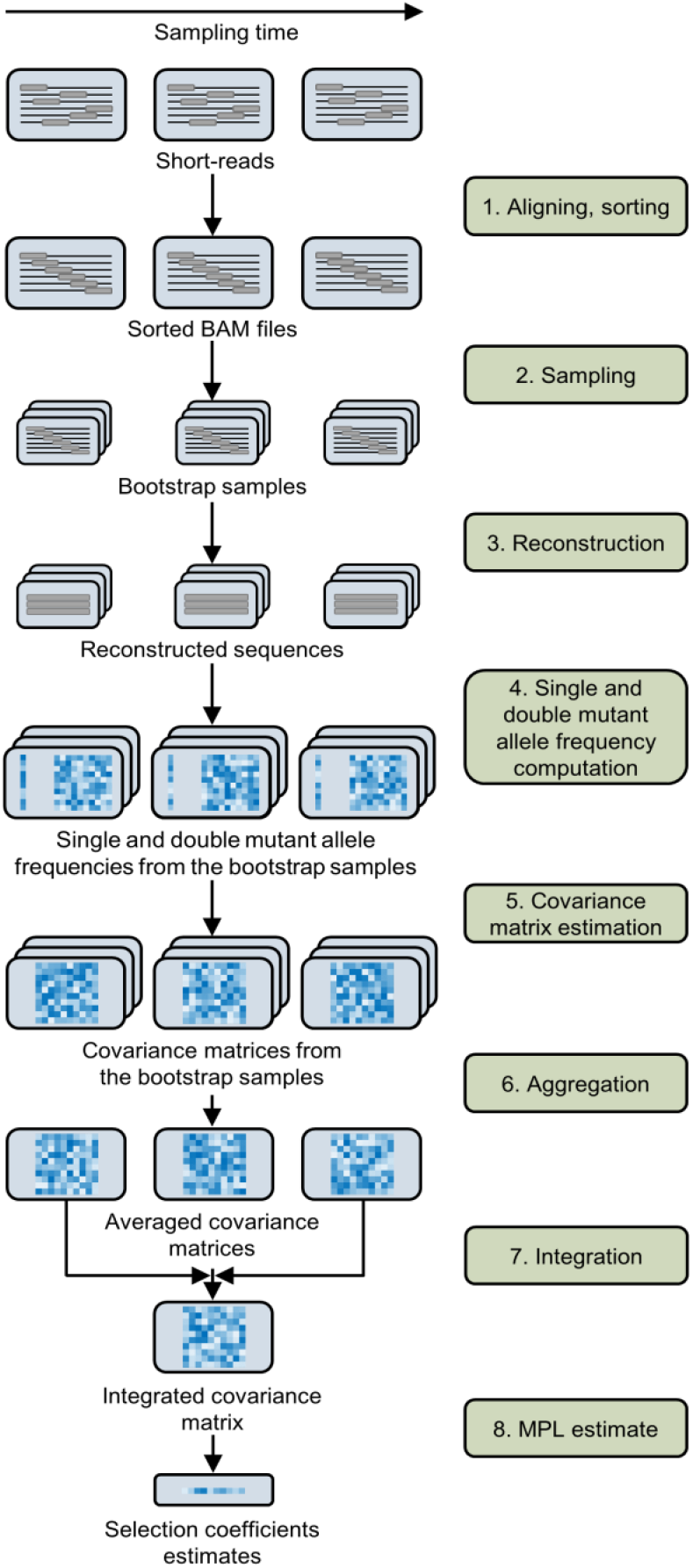
Illustration of the MPL-R pipeline. 1. Short reads from a longitudinal experiment are aligned to a reference sequence and sorted to obtain sorted BAM files. 2. Bootstrap samples are generated from the sorted BAM files by random sampling with replacement. 3. Population reconstruction is performed on the bootstrap samples to obtain a set of full-length sequences and their corresponding frequencies in the population. 4. For each sampled timepoint, the single and double mutant allele frequencies are obtained from the reconstructed sequences. 5. The covariance matrix of the mutant allele frequencies is computed for every bootstrap sample using the single and double mutant allele frequencies. 6. For each sampled time-point, the estimate of the covariance matrix of mutant allele frequencies of the actual population is obtained by averaging the covariance matrices of all *M* bootstrap samples. 7. The integrated covariance matrix (ICM), which quantifies the linkage effects, is obtained by scaling and summing these covariance matrices over all sampled time-points. 8. Accounting for linkage allows us to obtain accurate estimates of selection coefficients. The mutant allele frequencies obtained in Step 4 are also aggregated according to Equation 2 and used in Step 8, though not depicted here for simplicity.

### 2.2 Read aligning and sorting

The input to the pipeline consists of short-read data sampled at *K* + 1 time-points *t*_*k*_, for *k* ∈ {0, …, *K*} from an evolving population. For each time-point, the reads are aligned against a user-supplied reference sequence using BWA-MEM (Li, 2013) and the resulting *K* BAM files are sorted using samtools (Li *et al*., 2009).

### 2.3 Bagging and population reconstruction

The next step performs population reconstruction from short-read data. This is a computation and memory intensive procedure that presents a challenge, particularly as the number of reads becomes large. For efficient reconstruction, we pursue a bagging approach. We generate *M* bootstrap samples from each sorted BAM file such that each sample comprises of only a fraction of the total reads in the data (by random sampling with replacement). We then perform reconstruction using Quasirecomb on each bootstrap sample in parallel to obtain a probable set of full-length sequences constituting the population from which the short-read data is sampled. This approach is memory efficient (Breiman, 1999), while facilitating parallel processing to reduce the wall clock time required for the reconstruction process.

### 2.4 Estimation

The estimation step involves the calculation of a covariance matrix of mutant allele frequencies from each of the *M* reconstructed populations, each composed of *N* sequences of length *L*. The *L* × *L* covariance matrix *C*^*m*^(*t*_*k*_) of the mutant allele frequencies of the *m*-th bootstrap sample has (*i, j*)-th entry

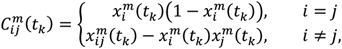

where 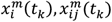 are the single mutant allele frequency and the double mutant allele frequency of the *m*-th bootstrap sample at locus *i* and locus-pair (*i, j*) at time *t*_*k*_ respectively. The covariance matrix of mutant allele frequencies, at time *t*_*k*_, *C*(*t*_*k*_), is then obtained by averaging the covariance matrices of all *M* bootstraps (the aggregating step of bagging), i.e.,

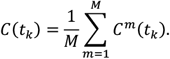

Scaling and summing the covariance matrices of mutant allele frequencies over the first *K* sampled time-points yields the integrated covariance matrix (ICM), *C*_int_, of size *L* × *L*, given as

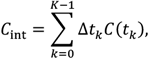

where Δ*t*_*k*_ = *t*_*k*+1_ − *t*_*k*_. The ICM captures linkage effects in the data over the course of evolution and is used to obtain the estimates of the selection coefficients, ŝ_*i*_, given by (Sohail *et al*., 2021)

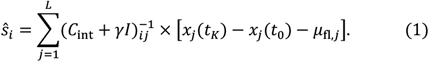

Here, *I* is an *L* × *L* identity matrix, *x*_*j*_(*t*_0_), *x*_*j*_(*t*_*K*_) are the single mutant allele frequencies at the first and the last sampled time-points of the *j*-th locus, respectively, and *γ* is a regularization parameter on the selection coefficients. Mathematically, *γ* = 1/(*Nσ*^2^) where *N* is the population size and *σ*^2^ is the variance of the Gaussian prior distribution for the selection coefficients (see Sohail *et al*., 2021 for more details). In practice, one does not need to set the value of *σ*^2^ as it is already incorporated into the inference when the value of *γ* is chosen (Simulated data). The regularization parameter ensures that the ICM, *C*_int_, is invertible. Regularization also suppresses the inference of large selection coefficients that are not well-supported by data. Moreover, *μ*_fl,*j*_ is the integrated mutational flux, representing the net change in allele frequencies due to random mutations. It is given by

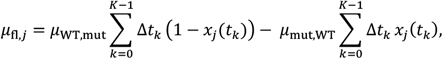

where *μ*_WT,mut_ and *μ*_mut,WT_ are the mutation probabilities of the WT allele to mutant allele and mutant allele to WT allele respectively. The single mutant allele frequencies of the *j*-th locus at time *t*_*k*_, *x*_*j*_(*t*_*k*_), are obtained by averaging the mutant allele frequencies of all *M* bootstrap samples,

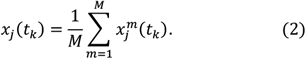

The number of polymorphic loci is smaller than the sequence length in real datasets. As only the polymorphic loci contribute towards parameter estimation, a practical methodology for parameter inference is to extract a submatrix and subvector corresponding to the polymorphic loci from the ICM and the numerator term in **Equation 1** respectively. This approach is faster with less storage requirements. In the pipeline, the ICM and the numerator term in **Equation 1** are prefiltered based on loci which have mutant alleles appearing at frequencies of ≥ 1% and inference is performed on this subset of loci. The mutant allele frequency at a locus is defined as the ratio between the number of reads carrying the mutation at the locus and the total number of reads covering the locus.

### 2.5 Simulated data

We simulated a Wright-Fisher model with selection and mutation. We considered a population of *N* = 1000 individuals, each represented by a *L* = 500 length binary string 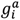, with zero representing the WT allele and one representing the mutant allele at locus *i* in sequence *a*. We assumed an additive fitness model with the total fitness of sequence *a* given by

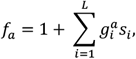

where *s*_*i*_ is the selection coefficient of the mutant allele at locus *i*. The forward and backward mutation probabilities were assumed to be equal for simplicity and were set to *μ* = 10^−4^ per locus per generation. Alleles at 20/20/460 loci were beneficial/deleterious/neutral respectively. The beneficial and deleterious mutant alleles had the same magnitude of selection coefficients, but they had opposite signs, while the selection coefficients of neutral alleles had a magnitude of zero. We simulated three structures of the underlying fitness landscape: “block” (Supplementary **Fig. S1A**), “comb” (Supplementary **Fig. S1B**), and “random” (Supplementary **Fig. S1C**). Unless otherwise stated, the underlying fitness landscape used the “comb-like” structure. We generated five sets of data, each with a different value of the strength of selection in {0.005, 0.010, 0.025, 0.050, 0.075}, two sets with the probability of recombination 10^−6^ and 10^−5^, two sets with *L* = 1500, *L* = 3000, and four sets with the number of founder sequences (sequences present in the population at the initial time-point) in the range 1 to 4. The founder sequences were randomly generated with the constraint that all founder sequences were within a Hamming distance of 10 to each other. We used 400 generations for inference with the population sampled at every Δ*t* = 50 generations.

We simulated short-read sequencing of the population using the ART simulator (Huang *et al*., 2012). At each time-point of the sets with *L* = 500, we generated *R* = 15,000 single-end reads of 150bp using the error profile of the Illumina HiSeq 2500 platform, giving a coverage of 4500X. We generated *R* = 45,000 and *R* = 90,000 reads at each time-point for the sets with *L* = 1,500 and *L* = 3,000 to ensure consistency of coverage. A total of 100 Monte Carlo runs were generated for each data set.

For inference, we generated *M* = 3 bootstrap samples by sampling (with replacement) from each sorted BAM file, with each bootstrap sample containing a third as many reads, i.e., *R*_*b*_ = *R*/3 = 5,000, 15,000, 30,000, for *L* = 500, 1,500, 3,000, as contained in the sorted BAM file. This amounted to a coverage of 1500X. The regularization parameter was set to *γ* = 10; though our results were insensitive to this choice, with MPL-R showing similar performance for a wide range of regularization values (Supplementary **Fig. S2**). We measured the performance of MPL-R to detect beneficial mutant alleles from the rest (neutral/deleterious) and deleterious mutant alleles from the rest (neutral/beneficial) in terms of the area under the receiver operating characteristic curve (AUROC). The AUROC was calculated on selection coefficients corresponding to mutant alleles that appeared at frequencies of ≥ 1%. On average, the mutant allele frequency trajectories of 14 of the 20 deleterious mutations crossed this threshold for the single-founder dataset with *L* = 500, *s* = 0.050, and no recombination.

### 2.6 HIV-1 data

We analyzed a real dataset of HIV-1 in vivo evolution to show the ability of our proposed pipeline to accurately infer selection from short-read data. As a benchmark, we used the analysis of the half-genome length time-series HIV-1 sequence data performed in Sohail *et al*., 2021. The HIV-1 dataset and its pre-processing were explained in detail in Sohail *et al*., 2021. Briefly, the dataset comprised Sanger-sequenced data from 13 HIV-1 infected patients sampled at different time-points as part of the Center for HIV/AIDS Vaccine Immunology (CHAVI 001) and Centre for the AIDS Programme of Research in South Africa (CAPRISA 002) studies in the United States, Malawi and South Africa. Sequence data was downloaded from Los Alamos National Laboratory HIV Sequence Database (www.hiv.lanl.gov). The sampling times and number of sequences at each time-point are described in Supplementary Table 1 in Sohail *et al*., 2021. The population size *N*, was not explicitly assumed as it was already incorporated into the value of *γ*. Mutation probabilities for all nucleotide transitions were obtained from Figure 1C in Zanini *et al*., 2017 and mutation probabilities to and from gap states were assumed to be 1 × 10^−9^ per locus per day following Sohail *et al*., 2021. These values have been summarized in Supplementary **Table S1**. We analyzed the group specific antigen (gag) protein to demonstrate the functionality of MPL-R.

We simulated sequencing of the gag protein (length 1,500bp) using the ART simulator with the error profile of the Illumina HiSeq 2500 to generate single-end reads of length 150bp. For ease of exposition, we set the sequencing parameters to 15 reads per sequence and 10,000 sequences resulting in *R* = 150,000 reads and an average coverage of 13,500X. After aligning and sorting the reads, we obtained *M* = 3 bootstrap samples for each time-point by sampling with replacement keeping the bootstrap sample size *R*_*b*_ = *R*/3 = 50,000.

Population reconstruction was performed as explained previously. The reconstructed sequences from each bootstrap sample were supplied to the MPL-R pipeline, preprocessed, and analyzed using the same parameter values used in Sohail *et al*., 2021 to obtain the ICM and the numerator term from **Equation 1**. The estimates of the ICM and the numerator term were averaged over all bootstrap samples, and the estimates of the selection coefficients obtained by the utilization of **Equation 1**. We also obtained the estimates of the selection coefficients from the full-length gag sequences via MPL for benchmarking.

The performance of MPL-R was compared against MPL via the proportion of biologically important mutations (those residing in CD8+ T cell epitopes, or nonsynonymous reversions outside CD8+ T cell epitopes, or nonsynonymous reversions within CD8+ T cell epitopes) in the top 5% strongest selection coefficients. Statistical significance of the proportions of biologically important mutations was computed via fold enrichment, which quantifies the likelihood of obtaining a particular proportion by pure chance. Mathematically, if *n*_obs_ mutations of a particular property are observed in a category with *N*_cat_ mutations, and there are *n*_tot_ mutations of that particular property in the total *N*_tot_ mutations, then the ratio (*n*_obs_/*N*_cat_)/(*n*_tot_/*N*_tot_) specifies the fold enrichment. The greater the value of fold enrichment relative to 1, the higher the confidence that *n*_obs_ is not observed by chance. Two-sided Fisher’s exact test was used to calculate the p-values of all values of fold-enrichment.

### 2.7 SARS-CoV-2 data

To demonstrate the working of the pipeline on another dataset, we performed inference on a real dataset of SARS-CoV-2 that was evolving in an immunocompromised patient infected with HIV (Velasquez-Reyes *et al*., 2025). The dataset consisted of 6 samples of 70bp paired-end reads sequenced via the Illumina NextSeq 500 platform from 312 to 776 days since the estimated date of SARS-CoV-2 infection. The average number of reads *R*, was 2,786,519 with a standard deviation of 1,008,062, which translates to a coverage of 4,658X with a standard deviation of 1,242X. The receptor-binding domain (RBD) of the Spike glycoprotein was chosen for analysis as it harbors biologically important mutations associated with antibody escape (Focosi *et al*. 2021, Tang *et al*. 2023, Cao *et al*. 2023). The coordinates of the RBD were obtained from Barnes *et al*., 2020. A plot of the single mutant allele frequencies (Supplementary **Fig. S3**) reported multiple simultaneous mutations across the genome and the RBD, which indicated linkage, and suggested that linkage-aware analysis could be informative. The reads were aligned against the SARS-CoV-2 reference sequence (GenBank accession no. MN908947.3) and sorted. *M* = 3 bootstrap samples were obtained at each time-point consisting of an average of *R*_*b*_ = *R*/3 reads as described before. Population reconstruction was performed between loci 22,544 to 23,161 to obtain full-length sequences of length 618bp of the RBD. Selection coefficients were estimated via a binary model of WT/mutant as described before. The mutation rate used in estimation was obtained from Velasquez-Reyes *et al*., 2025 and symmetric mutation rates were assumed between the WT allele to mutant allele and mutant allele to WT allele. The value of the regularization parameter was taken as *γ* = 10. Estimation was performed on only those loci where the single mutant allele frequency was ≥ 1%. Synonymous and non-synonymous mutations were classified via sc2calc (van Zwetselaar, 2021). After estimation, the performance of MPL-R was demonstrated by its ability to correctly identify known beneficial mutations in the RBD.

## 3 Results

### 3.1 Classification performance of MPL-R

The proposed computational pipeline MPL-R infers selection from shortread time series data. Initially, we tested the pipeline on simulated timeseries data (Methods). For comparison, we also evaluated two alternative schemes which did not require population reconstruction. The first scheme, MPL (banded), used the double mutant allele frequencies directly observable from the short-read data (i.e., the intra-read double mutant allele frequencies) to compute the covariance matrices and set the covariance matrix entries corresponding to the inter-read allele pairs to zero. The second scheme, termed MPL (identity covariance), used only the single mutant allele frequency information, and considered the loci to be under independent evolution. For benchmarking, we used MPL (Sohail *et al*., 2021) on the ground truth full-length sequences, with access to complete knowledge of the single and double mutant allele frequencies.

We considered a system with 500 loci that had simultaneously polymorphic loci spanning the entire length of the sequence. Analysis of the simulated data showed that only around 10% of pairs of polymorphic loci lay within one read length (150 bp) of each other; that is, most loci contributing to linkage effects are out-of-read polymorphic loci not directly observable with short-read data (**Fig. 2A**). We performed analysis using the schemes described and obtained estimates of the selection coefficients. Our results showed a high positive correlation (*ρ* = 0.72, p-value < 10^−100^) between the selection coefficients estimated by the proposed MPL-R method and those estimated by the benchmark MPL method (**Fig. 2B**). We also observed that the classification performance of the MPL-R method was close to that of the MPL method and was better than MPL (identity covariance) and MPL (banded) (**Fig. 2C**). The reduced performance of MPL-R as compared to MPL is because inference from shortread data has several sources of errors (sequencing errors, non-uniform sequencing coverage, and population reconstruction errors) not encountered in case of MPL (which assumes complete and accurate knowledge of full-length sequences). MPL (identity covariance) underperforms because it does not explicitly account for genetic linkage effects; thus, it over/underestimates selection coefficients of mutant alleles evolving under genetic linkage (Supplementary **Fig. S4**). Specifically, MPL (identity covariance) overestimates the selection coefficients of neutral alleles hitchhiking on beneficial ones and underestimates the magnitudes of the selection coefficients of mutant alleles evolving under clonal interference or negative selection. Incorporating the inferred inter-read covariances allows MPL-R to account for genetic linkage information and obtain more accurate selection coefficient estimates. This performance difference becomes more significant for selection coefficients with larger magnitudes (Supplementary **Fig. S5**), due to the stronger linkage effects in these cases. A more detailed explanation of the performance difference between MPL-R and MPL (identity covariance) is given in the Supplementary Information in Section SI.1.

**Fig. 2.**
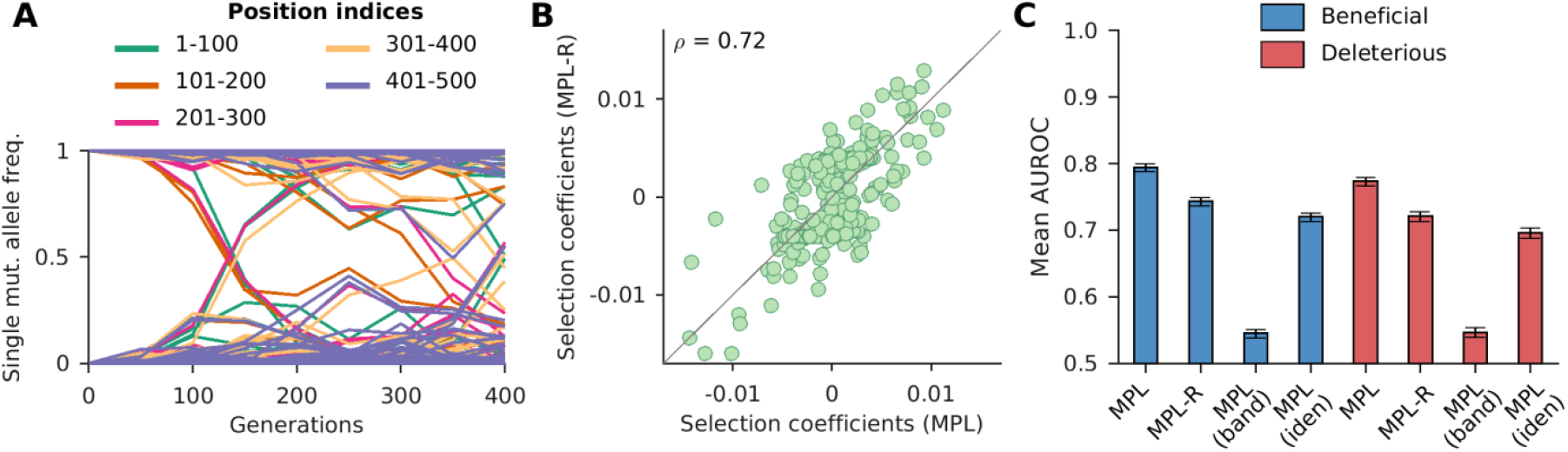
MPL-R has good classification performance. **(A)** Plot of single mutant allele frequency trajectories indicates the presence of polymorphic alleles across the entire length of the sequence. **(B**) Selection coefficients estimated from full-length sequences (i.e., all double mutant allele frequencies known) and those estimated from reconstructed double mutant allele frequencies were highly correlated (Pearson correlation coefficient *ρ* = 0.72 with a p-value < 10^−100^). Results shown here are for a typical Monte Carlo run. **(C)** The mean AUROC performance of distinguishing beneficial mutant alleles from the rest (deleterious and neutral), and deleterious mutant alleles from the rest (neutral and beneficial) of the proposed method (MPL-R) is compared against three methods: MPL, MPL (banded), and MPL (identity covariance). Results are shown for 100 Monte Carlo runs. Each Monte Carlo run consisted of evolving populations of *N* = 1000 individuals of *L* = 500 bi-allelic (WT and mutant) loci, with equal forward and backward mutation probabilities set to *µ* = 10^−4^ per locus per generation. Alleles at 20/20/460 loci were beneficial/deleterious/neutral with selection coefficients +0.025/-0.025/0 respectively. The fitness landscape had a repeating comb-like structure shown in Supplementary **Fig. S1B**. One founder sequence was used to generate each population, which was allowed to evolve for 400 generations and sampled at Δ*t* = 50 generations. Results for MPL-R are with *M* = 3 bootstrap samples each of size *R*_*b*_ = *R*/3 reads (coverage 1500X), where *R* is the total number of reads available at each sampled time-point. The error bars indicate the standard error of the mean.

Surprisingly, we observe that the MPL (banded) scheme, which attempts to account for within-read linkage effects while ignoring out-of-read effects, is outperformed by the MPL (identity covariance) scheme which ignores linkage effects completely. This suggests that the naïve approach of applying the MPL framework by substituting observable (within-read) double mutant allele frequencies and setting all unobservable double mutant allele frequencies to the product of the single mutant allele frequencies, is not a desirable approach. This has been discussed in detail in the Supplementary Information in Section SI.2. Briefly, while the ICM in MPL (banded) captures more information from the ICM obtained using the full-length sequences compared to the ICM in MPL (identity covariance), the same is not true for the respective inverse ICMs (Supplementary **Fig. S6**). Because of the inverse ICM term in **Equation 1**, MPL (identity covariance) outperforms MPL (banded). This result highlights the need for reconstructing full-length sequences from short-read data to infer unobserved inter-read covariance values, as incomplete linkage information obtained from short-read data (only using intra-read covariance information) is not sufficient for accurate inference of selection coefficients. The reconstruction step of MPL-R (Step 3 in **Fig. 1**) inferred the inter-read double mutant allele frequencies. These, in turn, were used to compute the ICM (Methods), which accounts for the genetic linkage present in the data. Our results showed that the proposed MPL-R method was able to accurately account for linkage effects across the length of the sequence, faithfully inferring ICM entries for both the intra and inter-read pairs (**Fig. 3**). Further simulations showed that the performance of MPL-R was robust to the level of diversity in the data (**Fig. 4A** and Supplementary **Fig. S7A**), the distribution of fitness effects across the length of the sequence (**Fig. 4B** and Supplementary **Fig. S7B**), the level of recombination (**Fig. 4C** and Supplementary **Fig. S7C**), and sequencing coverage (**Fig. 4D** and Supplementary **Fig. S7D**). The performance changed with changes in selection strength (**Fig. 4E** and Supplementary **Fig. S7E**), and the length of the genome (**Fig. 4F** and Supplementary **Fig. S7F**). The degradation of performance at low values of the strength of selection in **Fig. 4E** and Supplementary **Fig. S7E** is because very weak selection becomes hard to differentiate from neutral evolution, as the fluctuations in the mutant allele frequencies due to selection and random genetic drift become challenging to distinguish. The degradation associated with the increase in the length of the genome (**Fig. 4F** and Supplementary **Fig. S7F**) is explained by the fact that more linkage information becomes unobservable as the genome length increases relative to the read length. Population reconstruction and consequently inference of the unobservable values in the ICM become more difficult for larger sequences observed with a fixed read length. Supplementary **Fig. S8** demonstrates the degradation in the estimates of the ICM with the increase in the length of the genome while keeping the read length constant.

**Fig. 3.**
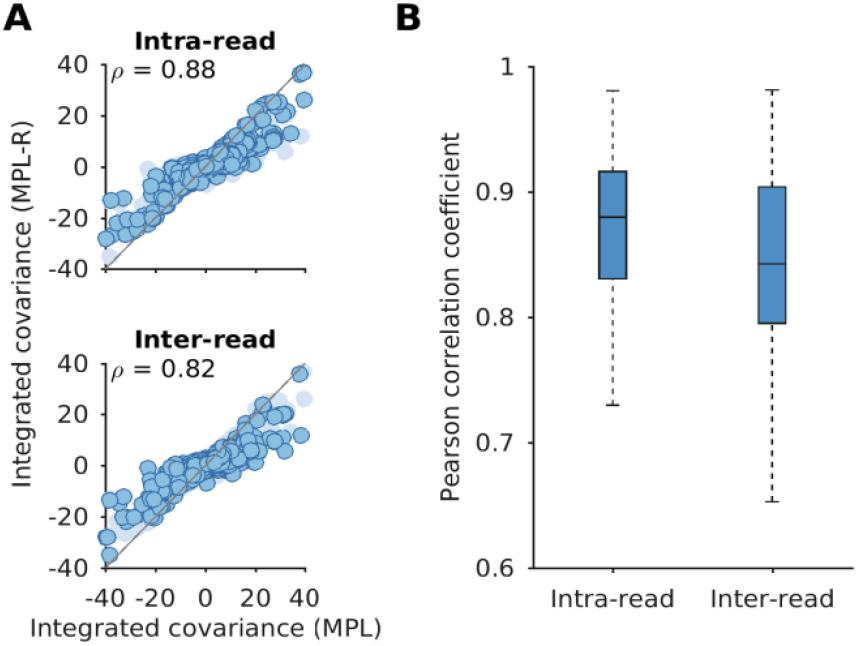
MPL-R recovers linkage effects across the entire sequence. The entries of the ICM calculated from the reconstructed double mutant allele frequencies exhibit high correlation with the entries of the ICM calculated from the double mutant allele frequencies obtained from the full-length sequences. **(A)** Intra-read covariance values (*top* panel) show good correlation (Pearson correlation coefficient *ρ* = 0.88 with p-value < 10^−100^). The interread covariance values (*bottom* panel) are highly correlated as well which demonstrate the ability of the method to infer linkage patterns not observable in individual short-reads (Pearson correlation coefficient *ρ* = 0.82 with p-value < 10^−100^). In each panel, the dark markers indicate the entries of the ICM in that category, and the light markers denote the rest of the entries. Results are shown here for a typical Monte Carlo run. **(B)** The summary statistics of the Pearson correlation coefficient for inter and intra-read covariance values indicate a consistent trend across 100 Monte Carlo runs. Each Monte Carlo run consisted of evolving populations of *N* = 1000 individuals of *L* = 500 bi-allelic (WT and mutant) loci, with equal forward and backward mutation probabilities set to *µ* = 10^−4^ per locus per generation. Alleles at 20/20/460 loci were beneficial/deleterious/neutral with selection coefficients +0.025/-0.025/0 respectively. The fitness landscape had a repeating comb-like structure shown in Supplementary **Fig. S1B**. One founder sequence was used to generate each population, which was allowed to evolve for 400 generations and sampled at Δ*t* = 50 generations. Results are for MPL-R with *M* = 3 bootstrap samples each of size *R*_*b*_ = *R*/3 reads (coverage 1500X), where *R* is the total number of reads available at each sampled timepoint.

**Fig. 4.**
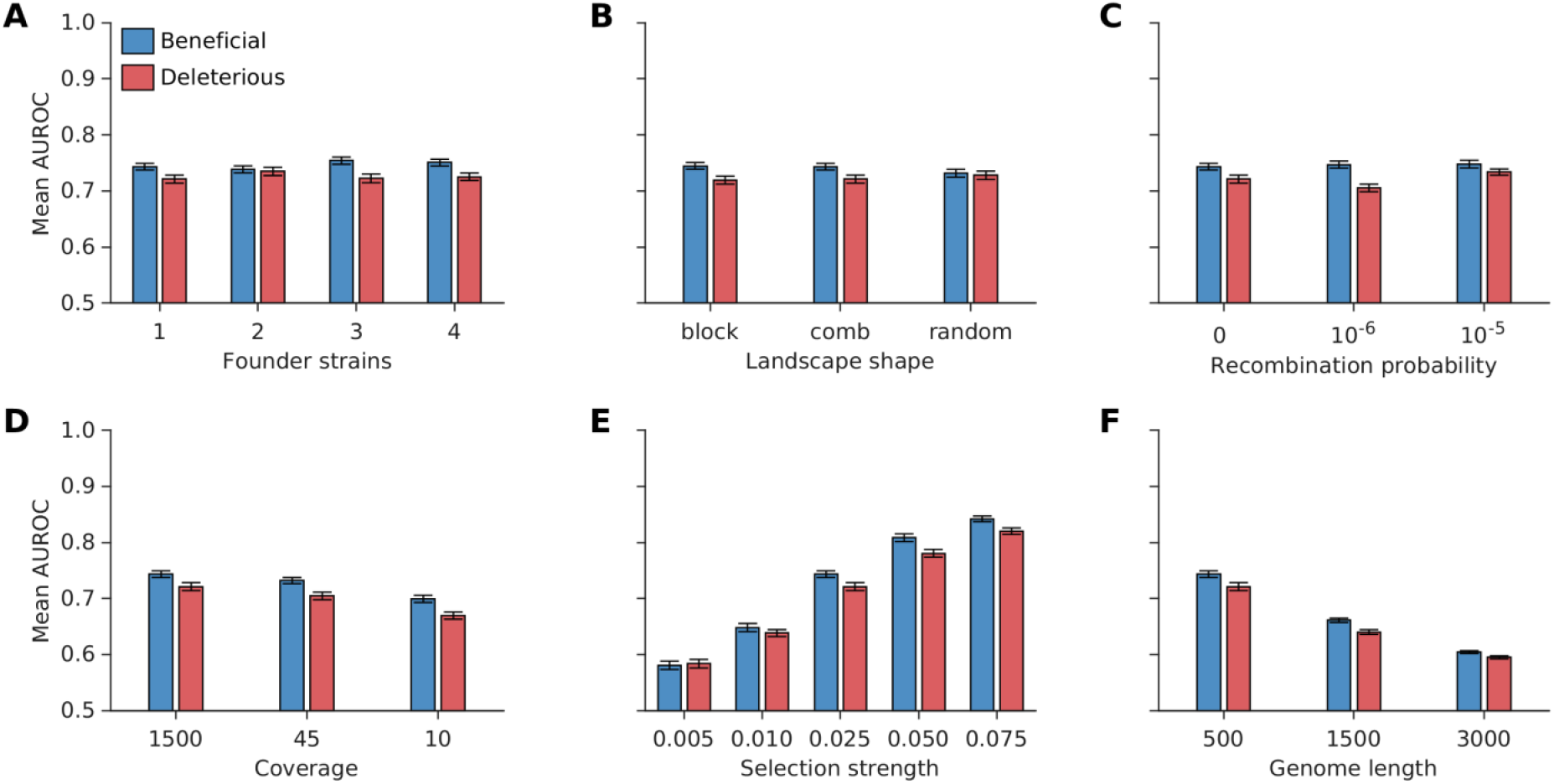
Robustness of MPL-R to changes in diversity in data, distribution of fitness effect across the length of the sequence, recombination probability, strength of selection, length of the genome, and coverage. The classification performance of MPL-R quantified as the mean AUROC remained largely unaffected for **(A)** increase in population diversity, controlled here by varying the number of founder strains (frequency of each founder ≥ 10%), **(B)** various structures of the underlying fitness landscape, **(C)** increase in recombination probability, and **(D)** sequencing coverage. The classification performance was affected by **(E)** decrease in selection strength, and **(F)** increase in the length of the genome. The structure of the data sets “comb”, “block”, and “random” is presented in Supplementary **Fig. S1**. Results are shown for 100 Monte Carlo runs. Each Monte Carlo run consisted of evolving populations of *N* = 1000 individuals of *L* = 500 bi-allelic (WT and mutant) loci, with equal forward and backward mutation probabilities set to *µ* = 10^−4^ per locus per generation. Unless mentioned otherwise, alleles at 20/20/460 loci were beneficial/deleterious/neutral with selection coefficients +0.025/-0.025/0 respectively. Unless mentioned otherwise, the fitness landscape had a repeating comb-like structure shown in Supplementary **Fig. S1B**. Unless mentioned otherwise, one founder sequence was used to generate each population, which was allowed to evolve for 400 generations and sampled at Δ*t* = 50 generations. Results are for MPL-R with *M* = 3 bootstrap samples each of size *R*_*b*_ = *R*/3 reads (coverage 1500X), where *R* is the total number of reads available at each sampled time-point. The error bars indicate the standard error of the mean.

### 3.2 Efficient reconstruction

The reconstruction step (Step 3 in **Fig. 1**) of MPL-R is by far the most time-consuming. However, the results above show that this step is necessary to resolve the effects of genetic linkage. The bagging approach (see Methods) used by MPL-R provides an efficient way to complete this step by forming *M* bootstrap samples at each sampled time-point, where each bootstrap sample consists of a fraction of the total number of reads available at that time-point. These bootstrap samples can be reconstructed on separate processors in parallel. Our simulations showed that the bagging approach employed by MPL-R resulted in a similar classification performance (**Fig. 5A**), and improved wall clock time as compared to MPL-R (full population reconstruction)—a scheme that performs population reconstruction on all the reads available at a sampled time-point (**Fig. 5B**). The wall clock time of the reconstruction step decreased by a factor of approximately *R*/*R*_*b*_, where *R* is the total number of reads available at a sampled time-point and *R*_*b*_ is the number of reads in a bootstrap sample. Simulations showed that decreasing *R*_*b*_, for the same number of bootstrap samples *M*, resulted in a proportional decrease in the wall clock time of the reconstruction step without affecting the classification accuracy (Supplementary **Fig. S9**).

**Fig. 5.**
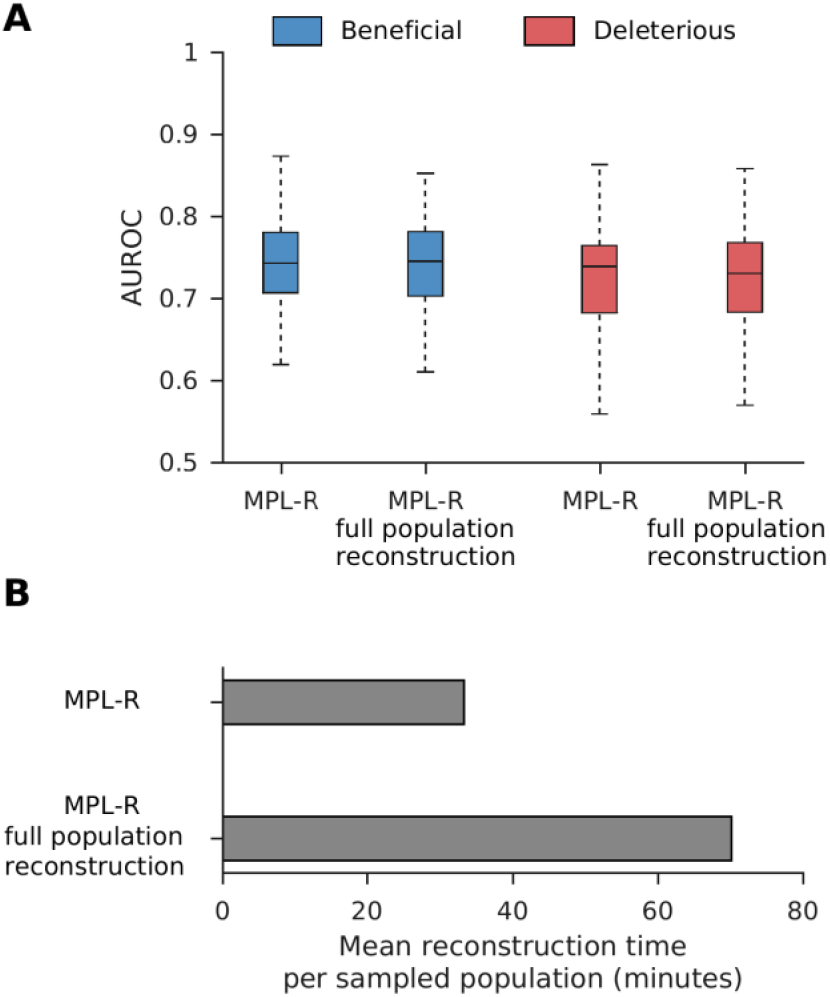
Bagging approach to reconstruction improves the wall clock time without affecting the classification performance. **(A)** Classification performance of the MPL-R method with bagging compared with a version that does not use bagging. Results of MPL-R are for *M* = 3 bootstrap samples each of size *R*_*b*_ = *R*/3 reads (coverage 1500X), where *R* is the total number of reads available at each sampled time-point used in MPL-R (full population reconstruction). **(B)** Average wall clock time of the reconstruction step of the proposed MPL-R method with bagging compared with MPL-R (full population reconstruction). Results are shown for 100 Monte Carlo runs. Each Monte Carlo run consisted of evolving populations of *N* = 1000 individuals of *L* = 500 bi-allelic (WT and mutant) loci, with equal forward and backward mutation probabilities set to *µ* = 10^−4^ per locus per generation. Alleles at 20/20/460 loci were beneficial/deleterious/neutral with selection coefficients +0.025/-0.025/0 respectively. The fitness landscape had a repeating comb-like structure shown in Supplementary **Fig. S1B**. One founder sequence was used to generate each population, which was allowed to evolve for 400 generations and sampled at Δ*t* = 50 generations.

### 3.3 Comparison with existing time-series methods

We sought to compare the proposed MPL-R method with other linkageaware methods in literature; namely IM (Illingworth, 2015), E&R (Terhorst *et al*., 2015), Evoracle (Shen *et al*., 2021), and Li and Barton, 2023. The method of Li and Barton, 2023 was the only linkage-aware method able to perform inference on the dataset, but it had poor classification performance, likely due to the large temporal sampling step used in the data. The other methods were hampered by impractically large wall clock times. For example, the IM method, the fastest of these methods, took ~4.5 hours for a single Monte Carlo run on an Intel Xeon 2.80 GHz CPU. Hence, a comparison of MPL-R’s performance with these methods was not feasible. MPL has been shown to outperform IM and E&R on smaller systems (Sohail *et al*., 2021), and since MPL-R has comparable performance as MPL, we expect MPL-R to outperform both IM and E&R. Other methods in the literature, e.g., LLS (Taus *et al*., 2017), FIT (Feder *et al*., 2014), WFABC (Foll *et al*., 2015), CLEAR (Iranmehr *et al*., 2017), and FITS (Zinger *et al*., 2019) use only the single mutant allele frequency information and do not seek to resolve linkage effects. Simulation results show that MPL-R uniformly outperformed these methods in terms of classification performance (**Fig. 6** and Supplementary **Fig. S10**). These methods did not require population reconstruction, and hence their wall clock times were in most cases significantly less than that of MPL-R. Nevertheless, the superior classification performance of MPL-R compensates for the additional computational requirements. Section SI.3 in the Supplementary Information presents further details of these alternative methods. It demonstrates empirically that MPL-R, even without compensating for genetic linkage, achieves a performance gain by explicitly accounting for mutations via the mutational flux term in **Equation 1** (**Fig. S11**).

**Fig. 6.**
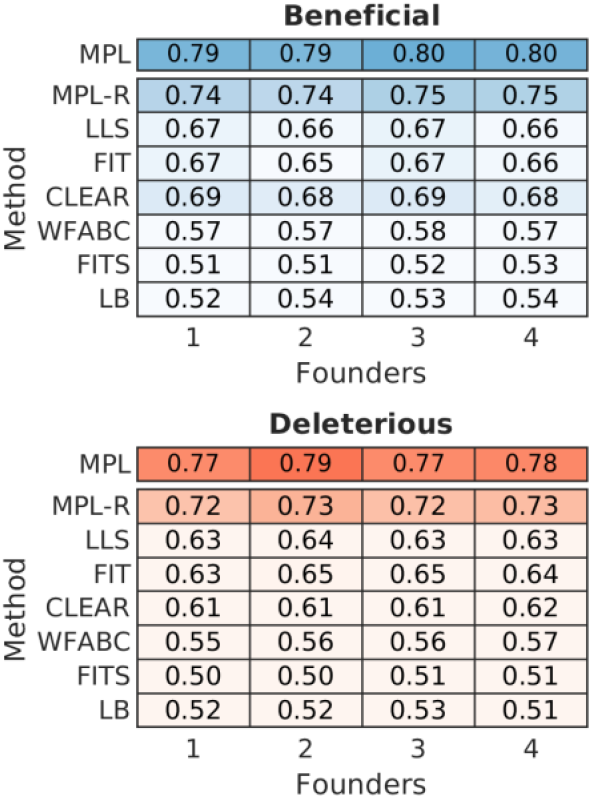
MPL-R outperforms available inference methods. The heatmap shows mean AUROC values of detecting beneficial alleles from the rest (deleterious and neutral) and detecting deleterious alleles from the rest (beneficial and neutral). A method with complete knowledge of all double mutant allele frequencies (MPL) is presented as a benchmark. Results of MPL-R are for *M* = 3 bootstrap samples each of size *R*_*b*_ = *R*/3 reads (coverage 1500X), where *R* is the total number of reads available at each sampled time-point. Results for the rest of the methods are based on the single mutant allele frequencies computed from the short reads obtained from the ART simulator. Results are shown for 100 Monte Carlo runs. Each Monte Carlo run consisted of evolving populations of *N* = 1000 individuals of *L* = 500 bi-allelic (WT and mutant) loci, with equal forward and backward mutation probabilities set to *µ* = 10^−4^ per locus per generation. Alleles at 20/20/460 loci were beneficial/deleterious/neutral with selection coefficients +0.025/-0.025/0 respectively. The fitness landscape had a repeating comb-like structure shown in Supplementary **Fig. S1B**. One founder sequence was used to generate each population, which was allowed to evolve for 400 generations and sampled at Δ*t* = 50 generations.

### 3.4 Application to HIV-1 patient data

Next, we analyzed longitudinal data of HIV-1 evolution in 13 patients (see Methods for details) using MPL-R. We identified the selection coefficients of mutant alleles associated with biological phenomenon such as CD8+ T cell escape, synonymous/nonsynonymous mutations, and reversions using methodology outlined in Sohail *et al*., 2021. **Fig. 7**. shows the composition of the top 5% mutations with the strongest selection coefficients estimated by MPL (*top-left* panel), MPL-R (*top-right* panel), and MPL (identity covariance) (*middle-left* panel), and the total composition of the mutations (*middle-right* panel). The percentages, enrichment, and p-values of CD8+ T cell escape mutations, nonsynonymous reversions outside CD8+ T cell epitopes, and nonsynonymous reversions within CD8+ T cell epitopes in the top 5% strongest mutations are reported in Supplementary **Table S2**. MPL-R and MPL both report higher fractions of CD8+ T cell escape mutations and lower fractions of nonsynonymous reversions outside CD8+ T cell epitopes as compared to MPL (identity covariance). Resolution of unobservable linkage information (*bottom* panel) enabled MPL-R to yield insights that were more in-line with those obtained via full-length sequences, which were in turn, had been shown to be consistent with known biological results (Sohail *et al*., 2021). In terms of timing performance, MPL-R was able to perform the analysis in a reasonable amount of time (Supplementary **Fig. S12**).

**Fig. 7.**
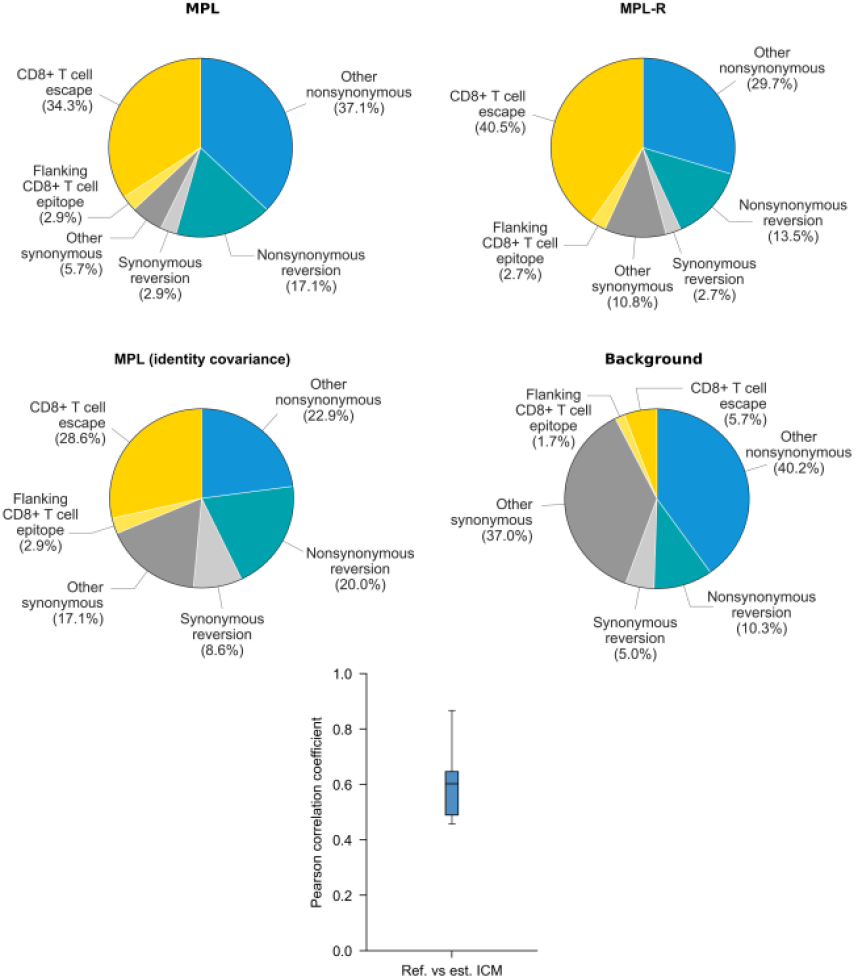
MPL-R correctly infers biological insights from intrahost HIV-1 patient data. The proportion of escape mutations and nonsynonymous reversions in the top 5% strongest selection coefficients identified by MPL-R (*top-right* panel) are similar to MPL (*top-left* panel). MPL (identity covariance) is not able to accurately reproduce the insights obtained via MPL (*middle-left* panel). The total composition of the mutations is shown as the background (*middle-right* panel). The ICMs estimated by MPL-R demonstrate moderate correlation with the ground truth ICMs (*bottom* panel).

### 3.5 Application to SARS-CoV-2 patient data

Next, to demonstrate MPL-R’s ability to account for linkage while inferring fitness effects of mutations from empirical short read data we analyzed intrapatient data of SARS-CoV-2 evolution (see Methods for details). We used LoFreq (Wilm *et al*., 2012) to count the number of non-synonymous mutations that exhibited at least 1% variation in the single mutant allele frequency. LoFreq reported 510 such nonsynonymous mutations across the entire genome, of which 26 were in the RBD. The number of mutations in the RBD was 2.6 times (two-sided Fisher’s exact test p-value 2.71× 10^−5^) higher than expected by random chance. An elevated number of mutations in the RBD suggested immune pressure on the virus. This was further supported by analysis of the dN/dS (ratio of non-synonymous and synonymous mutations), which evaluated to a mean value of 7.789±2.706 for the RBD. The selection coefficients corresponding to nonsynonymous mutations in the RBD, when inferred by MPL-R (which accounts for linkage effects) had higher magnitudes as compared to those inferred by MPL (identity covariance); a method that does not account for linkage effects (**Fig. 8**). Specifically, MPL-R inferred stronger positive selection coefficient estimates, classifying most mutations in the RBD as beneficial, while MPL (identity covariance) inferred relatively weaker estimates of selection coefficients, classifying most of these mutations as weakly beneficial or deleterious. We searched existing literature for all mutations in **Fig. 8** for which MPL-R and MPL (identity covariance) reported selection coefficients with opposite signs or differing by more than one order of magnitude. Supplementary **Table S3** shows that all RBD mutations classified as beneficial by MPL-R and underestimated by MPL (identity covariance) have been reported to be beneficial to the virus. The majority of these mutations benefit the virus by their contribution in antibody escape, whereas some mutations enhance infectivity or binding to the host (see Supplementary **Table S3** for details). The observation that MPL-R can identify more biologically-validated beneficial mutations as compared to MPL (identity covariance) demonstrates the advantage of the incorporation of linkage information.

**Fig. 8.**
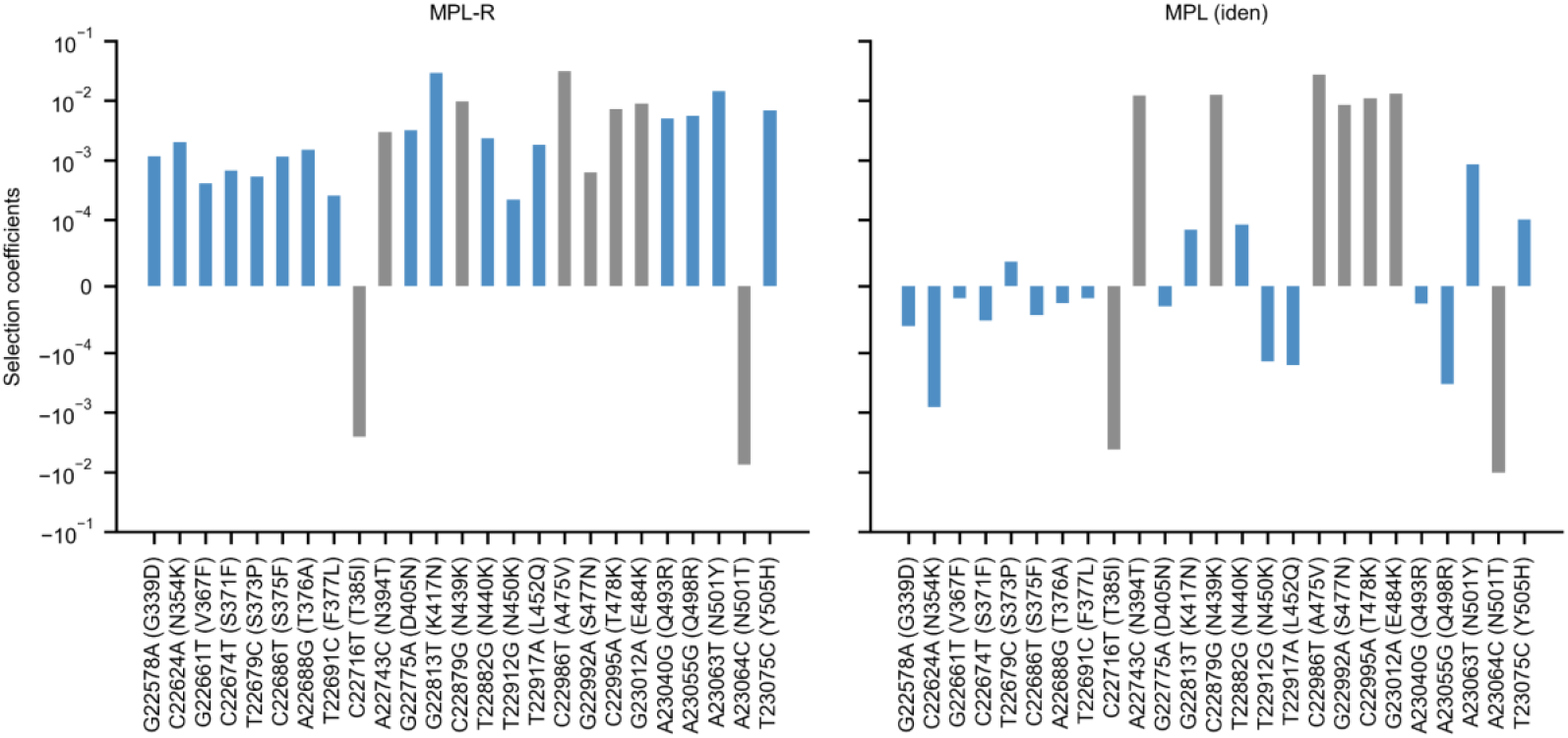
MPL-R identifies a higher number of beneficial mutations in the RBD. MPL-R identifies many known beneficial mutations (*left* panel), while MPL (identity covariance) infers several beneficial mutations as neutral or deleterious (*right* panel). Mutations where the estimates of MPL-R and MPL (identity covariance) differ by an order of magnitude or have opposite signs are highlighted in blue. Experimentally-reported phenotypic effects of the mutations are summarized in Supplementary **Table S3**.

## 4 Discussion

Short-read sequencing is a popular sequencing technology that is used extensively (Goodwin *et al*., 2016; Nafea *et al*., 2024). In this study, we have developed MPL-R, a computational pipeline for the inference of selection coefficients from short-read time-series data and demonstrated its accuracy over a range of parameters. Our analysis revealed novel insight into the importance of the off-diagonal ICM entries. We have shown that if the ICM entries corresponding to the unobservable inter-read allele-pairs are set to zero, the inference performance deteriorates substantially compared to models that assume independent evolution (Supplementary Information Section SI.2). We also presented an empirical result which demonstrates that incorporation of the mutational flux term assists in more accurate inference. This is an added advantage of MPL-based methods over methods that do not explicitly model mutation.

Any inference method based on population reconstruction is bound to face two key issues, which MPL-R addresses. First, the reconstruction time scales linearly with the number of reads, which implies that population reconstruction becomes increasingly difficult as sequencing platforms improve and read throughput increases. Here the engineering novelty of MPL-R comes into play. To our knowledge, MPL-R demonstrated for the first time the utility of a divide-and-conquer bagging approach of population reconstruction for inference of selection coefficients. MPL-R is designed to efficiently perform inference by making use of bagging, which ensures that the volume of reads does not significantly affect reconstruction time. MPL-R also distributes memory consumption over multiple reconstruction operations which can be useful when dealing with large datasets. The second issue addressed is that all reconstruction algorithms struggle with accurate population reconstruction, particularly when the underlying population contains multiple haplotypes or haplotypes with low frequencies (Posada-Cespedes *et al*., 2017). MPL-R works at the level of single and double mutant allele frequencies so is expected to stay immune to errors in higher-order statistics. Our simulations showed that the double mutant allele frequencies reconstructed by the pipeline were accurate (**Fig. 9** and Supplementary **Fig. S13**). The ICM entries computed from the reconstructed double mutant allele frequencies were accurate as well, as shown in **Fig. 3A**. This is sufficient for the purpose of inferring selection coefficients as the MPL framework only requires knowledge of single and double mutant allele frequencies, while errors in frequencies of higher-order tuples do not affect the accuracy of the estimated selection coefficients (Sohail *et al*., 2021). The higher-order frequencies may be important for more complex models, such as those incorporating epistasis (Sohail *et al*., 2022). Further work can explore the extent to which higher order frequencies are preserved by population reconstruction using Quasirecomb or other approaches. An initial analysis of triple mutant allele frequencies suggests that those obtained after reconstruction were strongly correlated with the triple mutant allele frequencies of the original population (Supplementary **Fig. S14**), though more extensive future investigation is warranted. In our tests, although the reconstruction was not perfect, Quasirecomb did a fairly good job of reconstructing the full-length sequences. A distribution of the mean Hamming distance between the reconstructed sequences and the closest sequence in the reference population had a median of around 3.5, which showed an average of around one erroneous bp per read-length (Supplementary **Fig. S15A**). The distribution of the mean Hamming distance between the reference sequences and the closest sequence in the reconstructed populations also had a median of around 3.5, indicating that all the reference sequences were reconstructed with few errors. The low value of the median Earth Mover’s distance (Huddleston *et al*., 2020) between the reference and reconstructed populations and between the reconstructed populations also suggested good reconstruction (Supplementary **Fig. S15B**).

**Fig. 9.**
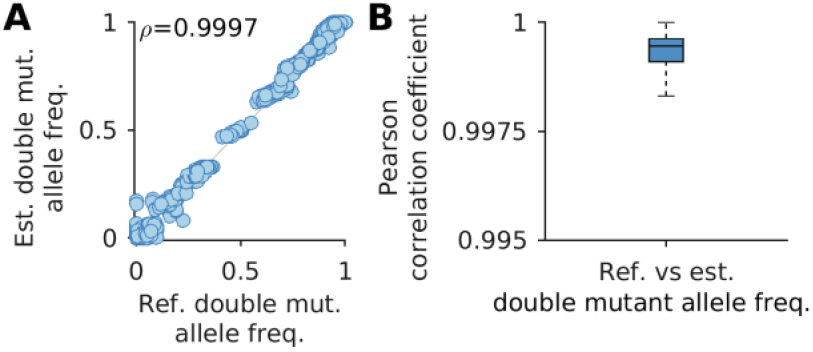
Reconstruction of double mutant allele frequencies is accurate. **(A)** Double mutant allele frequencies estimated after reconstruction show high correlation with the actual double mutant allele frequencies (Pearson correlation coefficient *ρ* = 0.9997 with p-value < 10^−100^). **(B)** The summary statistics of the Pearson correlation coefficient between the reference and estimated double mutant allele frequencies indicate a consistent trend across all Monte Carlo runs. Results are for MPL-R with *M* = 1 bootstrap sample of size *R*_*b*_ = *R*/3 reads (coverage 1500X), where *R* is the total number of reads available at each sampled time-point. Results are shown for 100 Monte Carlo runs. Each Monte Carlo run consisted of evolving populations of *N* = 1000 individuals of *L* = 500 bi-allelic (WT and mutant) loci, with equal forward and backward mutation probabilities set to *µ* = 10^−4^ per locus per generation. Alleles at 20/20/460 loci were beneficial/deleterious/neutral with selection coefficients +0.025/-0.025/0 respectively. The fitness landscape had a repeating comb-like structure shown in Supplementary **Fig. S1B**. One founder sequence was used to generate each population, which was allowed to evolve for 400 generations and sampled at Δ*t* = 50 generations.

The covariance matrices calculated from the reconstructed double mutant allele frequencies showed strong correlation with the covariance matrices calculated from the double mutant allele frequencies obtained from the full-length sequences (Supplementary **Fig. S16**). The initial time-points demonstrated weak correlation because the single mutant allele frequencies at these time-points were small (**Fig. 2A**), which resulted in covariance matrix entries with negligible magnitudes. At later time-points, the single mutant allele frequencies had larger magnitudes, which led to relatively larger covariance matrix entries, and stronger correlation.

MPL-R provides users with an easy-to-use bioinformatics pipeline to infer selection coefficients from short-read time-series data. The pipeline not only combines the features of its constituent tools but introduces the functionality of handling data transfer, intermediate processing, and function calls internally to minimize the workload of the end-user. The pipeline is flexible in terms of the components used and is scalable to the data sizes tested. MPL-R has been tested on data from a single sequencing platform, yet the reconstruction scheme based on Quasirecomb is general and can reconstruct data from several different sequencing platforms. The proposed MPL-R method offers a novel method to perform fitness inference of genetic time-series data and can be applied to study the complex evolutionary dynamics of populations of evolving microbes. Future work can explore fitness inference from long-read sequencing data and address the incorporation, testing, and optimization of software for long-read processing into the pipeline.

## 5 Software availability

The pipeline for MPL-R and example input files are accessible online at https://github.com/SMUAbdullah/paper-MPL-short-reads.

## 7 Acknowledgement

This work was carried out using the computational facilities of the Central High Performance Computing Cluster provided by the Information Technology Services Office and the computational systems of the Information Technology Office in the Department of Electronic & Computer Engineering at The Hong Kong University of Science and Technology. Simulations were also carried out on the High-Performance Computing Cluster provided by the Department of Electrical Engineering and CityU HK Burgundy High-Performance Computing system managed and provided by the Computing Services Centre at City University of Hong Kong.

## References

Alam, M.T. et al. (2011) Selective sweeps and genetic lineages of Plasmodium falciparum drug-resistant alleles in Ghana. J. Infect. Dis., 203, 220–227.

Barnes, C. O. et al. (2020) Structures of human antibodies bound to SARS-CoV-2 spike reveal common epitopes and recurrent features of antibodies. Cell, 182, 828–842.

Barroso-Batista, J. et al. (2020) Specific eco-evolutionary contexts in the mouse gut reveal Escherichia coli metabolic versatility. Curr. Biol., 30, 1049-1062.e7.

Van Den Bergh, B. et al. (2016) Frequency of antibiotic application drives rapid evolutionary adaptation of Escherichia coli persistence. Nat. Microbiol., 1, 1–7.

Bertels, F. et al. (2019) Parallel evolution of HIV-1 in a long-term experiment. Mol. Biol. Evol., 36, 2400–2414.

Bollback, J.P. et al. (2008) Estimation of 2 N es from temporal allele frequency data. Genetics, 179, 497–502.

Boshier, F.A.T. et al. (2022) Evolution of viral variants in remdesivir-treated and untreated SARS-CoV-2-infected pediatrics patients. J. Med. Virol., 94, 161–172.

Breiman, L. (1996) Bagging predictors. Mach. Learn., 24, 123–140.

Breiman, L. (1999) Pasting small votes for classification in large databases and on-line. Mach. Learn., 36, 85–103.

Buffalo, V. & Coop, G. (2019) The linked selection signature of rapid adaptation in temporal genomic data. Genetics, 213, 1007–1045.

Bull, R.A. et al. (2011) Sequential bottlenecks drive viral evolution in early acute hepatitis C virus infection. PLoS Pathog., 7, e1002243.

Cao, Y. et al. (2023). Imprinted SARS-CoV-2 humoral immunity induces convergent Omicron RBD evolution. Nature, 614, 521–529.

Charlesworth, B. (1994) The effect of background selection against deleterious mutations on weakly selected, linked variants. Genet. Res., 63, 213–227.

Charlesworth, B. et al. (1993) The effect of deleterious mutations on neutral molecular variation. Genetics, 134, 1289–1303.

Cheng, Q. et al. (2025) Selection estimation from genetic time-series data: Effects of limited sampling and genetic drift. Mol. Biol. Evol., msaf301.

Cordey, S. et al. (2010) Rhinovirus genome evolution during experimental human infection. PLoS One, 5, e10588.

van Dijk, E.L. et al. (2014) Ten years of next-generation sequencing technology. Trends Genet., 30, 418–426.

Elshire, R.J. et al. (2011) A robust, simple genotyping-by-sequencing (GBS) approach for high diversity species. PLoS One, 6, e19379.

Feder, A.F. et al. (2014) Identifying signatures of selection in genetic time series. Genetics, 196, 509–522.

Ferrer-Admetlla, A. et al. (2016) An approximate Markov Model for the Wright-Fisher diffusion and its application to time series data. Genetics, 203, 831–846.

Fischer, A. et al. (2014) High-definition reconstruction of clonal composition in cancer. Cell Rep., 7, 1740–1752.

Fisher, K.J. et al. (2018) Adaptive genome duplication affects patterns of molecular evolution in Saccharomyces cerevisiae. PLoS Genet., 14, e1007396.

Focosi, D. et al. (2021). Analysis of immune escape variants from antibody-based therapeutics against COVID-19: a systematic review. Int. J. Mol. Sci., 23, 29.

Foll, M. et al. (2015) WFABC: a Wright-Fisher ABC-based approach for inferring effective population sizes and selection coefficients from time-sampled data. Mol. Ecol. Resour., 15, 87–98.

Frenkel, E.M. et al. (2014) The fates of mutant lineages and the distribution of fitness effects of beneficial mutations in laboratory budding yeast populations. Genetics, 196, 1217–1226.

Gao, Y. and Barton, J.P. (2025) A binary trait model reveals the fitness effects of HIV-1 escape from T cell responses. Proc. Natl. Acad. Sci. U.S.A., 122, e2405379122.

Gerrish, P. and Lenski, R. (1998) The fate of competing beneficial mutations in an asexual population. Genetica, 102, 127–144.

Goodwin, S. et al. (2016) Coming of age: ten years of next-generation sequencing technologies. Nat. Rev. Genet., 17, 333–351.

Griffin, P.C. et al. (2017) Genomic trajectories to desiccation resistance: convergence and divergence among replicate selected Drosophila lines. Genetics, 205, 871–890.

He, Z. et al. (2020) Detecting and quantifying natural selection at two linked loci from time series data of allele frequencies with forward-in-time simulations. Genetics, 216, 521–541.

Hedrick, P.W. (1987) Gametic disequilibrium measures: proceed with caution. Genetics, 117, 331–341.

Hegreness, M. et al. (2006) An equivalence principle for the incorporation of favorable mutations in asexual populations. Science, 311, 1615–1617.

Huang, W. et al. (2012) ART: a next-generation sequencing read simulator. Bioinform., 28, 593–594.

Huddleston, J. et al. (2020) Integrating genotypes and phenotypes improves long-term forecasts of seasonal influenza A/H3N2 evolution. eLife, 9, 1–48.

Illingworth, C.J.R. (2015) Fitness inference from short-read data: withinhost evolution of a reassortant H5N1 influenza virus. Mol. Biol. Evol., 32, 3012–3026.

Imai, M. et al. (2012) Experimental adaptation of an influenza H5 HA confers respiratory droplet transmission to a reassortant H5 HA/H1N1 virus in ferrets. Nature, 486, 420–428.

Iranmehr, A. et al. (2017) CLEAR: Composition of likelihoods for evolve and resequence experiments. Genetics, 206, 1011–1023.

Ishaque, N. et al. (2018) Whole genome sequencing puts forward hypotheses on metastasis evolution and therapy in colorectal cancer. Nat. Commun., 9, 1–14.

Kemp, S.A. et al. (2021) SARS-CoV-2 evolution during treatment of chronic infection. Nature, 592, 277–282.

Lacerda, M. and Seoighe, C. (2014) Population genetics inference for longitudinally-sampled mutants under strong selection. Genetics, 198, 1237–1250.

Lang, G.I. et al. (2013) Pervasive genetic hitchhiking and clonal interference in forty evolving yeast populations. Nature, 500, 571–574.

Lee, B. et al. (2025) Inferring effects of mutations on SARS-CoV-2 transmission from genomic surveillance data. Nat. Commun., 16, 441.

Lee, K.T. et al. (2024) Benign polymorphisms in the BRCA genes with linkage disequilibrium is associated with cancer characteristics. Cancer Sci., 115, 3973–3985.

Li, H. (2013) Aligning sequence reads, clone sequences and assembly contigs with BWA-MEM. arXiv preprint 1303.3997.

Li, H. et al. (2009) The sequence alignment/map format and SAMtools. Bioinform., 25, 2078–2079.

Li, Y. and Barton, J.P. (2023) Estimating linkage disequilibrium and selection from allele frequency trajectories. Genetics, 223, iyac189.

Łuksza, M. and Lässig, M. (2014) A predictive fitness model for influenza. Nature, 507, 57–61.

Malaspinas, A. S. et al. (2012) Estimating allele age and selection coefficient from time-serial data. Genetics, 192, 599–607.

Mathieson, I. and McVean, G. (2013) Estimating selection coefficients in spatially structured populations from time series data of allele frequencies. Genetics, 193, 973–984.

McCrone, J.T. et al. (2018) Stochastic processes constrain the within and between host evolution of influenza virus. eLife, 7, e35962.

Metzker, M.L. (2010) Sequencing technologies-the next generation. Nat. Rev. Genet., 11, 31–46.

Nafea, A.M. et al. (2024) Application of next-generation sequencing to identify different pathogens. Front. Microbiol., 14, 1329330.

Orozco-Terwengel, P. et al. (2012) Adaptation of Drosophila to a novel laboratory environment reveals temporally heterogeneous trajectories of selected alleles. Mol. Ecol., 21, 4931–4941.

Ozsolak, F. and Milos, P.M. (2011) RNA sequencing: advances, challenges and opportunities. Nat. Rev. Genet., 12, 87–98.

Posada-Cespedes, S. et al. (2017) Recent advances in inferring viral diversity from high-throughput sequencing data. Virus Res., 239, 17–32.

Raynes, Y. et al. (2018) Sign of selection on mutation rate modifiers depends on population size. Proc. Natl. Acad. Sci. U.S.A., 115, 3422–3427.

Sanchez-Mazas, A. et al. (2017) The HLA-B landscape of Africa: signatures of pathogen-driven selection and molecular identification of candidate alleles to malaria protection. Mol. Ecol., 26, 6238–6252.

Schirmer, M. et al. (2014) Benchmarking of viral haplotype reconstruction programmes: an overview of the capacities and limitations of currently available programmes. Brief. Bioinform., 15, 431–442.

Schraiber, J.G. et al. (2016) Bayesian inference of natural selection from allele frequency time series. Genetics, 203, 493–511.

Setliff, I. et al. (2018) Multi-donor longitudinal antibody repertoire sequencing reveals the existence of public antibody clonotypes in HIV-1 infection. Cell Host Microbe, 23, 845–854.

Shen, M.W. et al. (2021) Reconstruction of evolving gene variants and fitness from short sequencing reads. Nat. Chem. Biol., 17, 1188–1198.

Smith, J.M. and Haigh, J. (1974) The hitch-hiking effect of a favourable gene. Genet. Res., 23, 23–35.

Sohail, M.S. et al. (2022) Inferring epistasis from genetic time-series data. Mol. Biol. Evol., 39, msac199.

Sohail, M.S. et al. (2021) MPL resolves genetic linkage in fitness inference from complex evolutionary histories. Nat. Biotechnol., 39, 472–479.

Strelkowa, N. and Lässig, M. (2012) Clonal interference in the evolution of influenza. Genetics, 192, 671–682.

Takeda, H. et al. (2017) Evolution of multi-drug resistant HCV clones from pre-existing resistant-associated variants during direct-acting antiviral therapy determined by third-generation sequencing. Sci. Rep., 7, 45605.

Tang, X. et al. (2023). Adaptive evolution of the spike protein in coronaviruses. Mol. Biol. Evol., 40, msad089.

Taus, T. et al. (2017) Quantifying selection with pool-seq time series data. Mol. Biol. Evol., 34, 3023–3034.

Terhorst, J. et al. (2015) Multi-locus analysis of genomic time series data from experimental evolution. PLoS Genet., 11, e1005069.

Topa, H. et al. (2015) Gaussian process test for high-throughput sequencing time series: application to experimental evolution. Bioinform., 31, 1762–1770.

Töpfer, A. et al. (2013) Probabilistic inference of viral quasispecies subject to recombination. J. Comput. Biol., 20, 113–123.

van Zwetselaar M. (2021) sc2calc - SARS-CoV-2 genome coordinate converter. https://github.com/zwets/sc2calc (accessed October 20, 2025).

Velasquez-Reyes, J. M. et al. (2025) Characterisation of a persistent SARS-CoV-2 infection lasting more than 750 days in a person living with HIV: a genomic analysis. The Lancet Microbe, 6.

Wilm, A. et al. (2012) LoFreq: a sequence-quality aware, ultra-sensitive variant caller for uncovering cell-population heterogeneity from high-throughput sequencing datasets. Nucleic Acids Res., 40, 11189–11201.

Xue, K.S. et al. (2018) Within-host evolution of human influenza virus. Trends Microbiol., 26, 781–793.

Zanini, F. et al. (2015) Population genomics of intrapatient HIV-1 evolution. eLife, 4, e11282.

Zanini, F. et al. (2017) In vivo mutation rates and the landscape of fitness costs of HIV-1. Virus Evol., 3, vex003.

Zeng H et al. (2024) Two fitness inference schemes compared using allele frequencies from 1,068,391 sequences sampled in the UK during the COVID-19 pandemic, Phys. Biol., 22, 016003.

Zinger, T. et al. (2019) Inferring population genetics parameters of evolving viruses using time-series data. Virus Evol., 5, vez011.

