## supplementary_information for "Linkage-aware inference of fitness from short-read time-series genomic data"

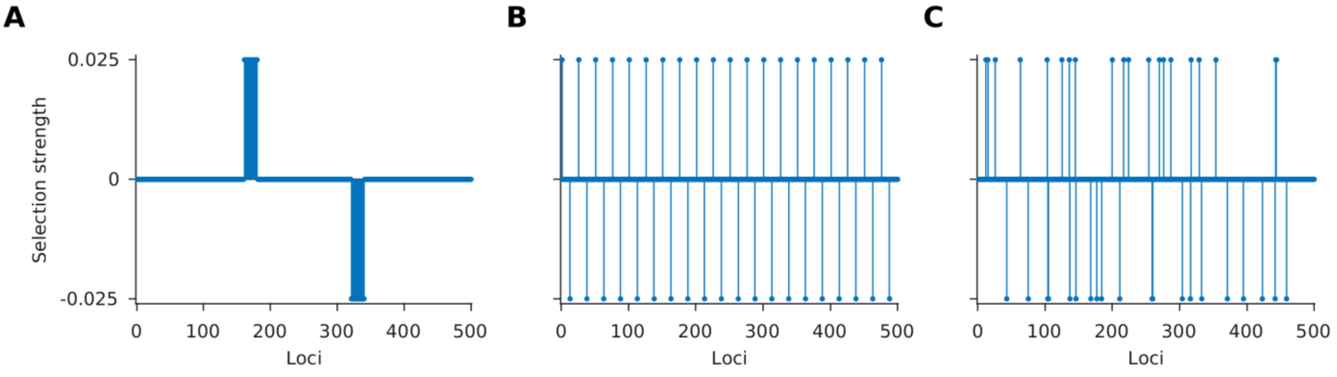


**Fig. S1. Structures of fitness landscape used in the simulations. (A)** The data set "block" contained a single block of beneficial alleles grouped together at loci 161 to 180, a block of deleterious alleles at the loci 321 to 340, and neutral alleles at all the remaining loci. **(B)** The data set "comb" contained a repeating structure of 1 beneficial-11 neutral-1 deleterious-12 neutral alleles. **(C)** The data set "random" had the beneficial, deleterious, and neutral alleles distributed randomly across the length of the sequence.

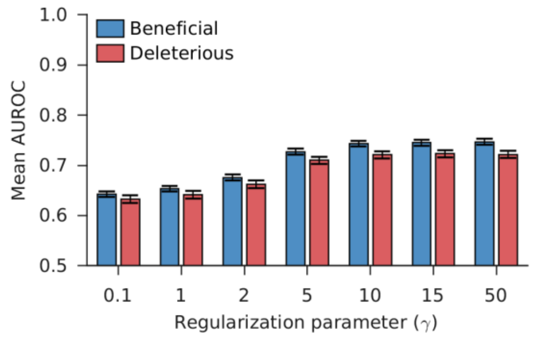


**Fig. S2. Exact value of the regularization parameter does not affect classification performance.** The classification performance quantified as the mean AUROC is not sensitive to the value of the regularization parameter for values between 5 and 50. Results are shown for 100 Monte Carlo runs. Each Monte Carlo run consisted of evolving populations of $N = 1000$ individuals of $L = 500$ bi-allelic (WT and mutant) loci, with equal forward and backward mutation probabilities set to $\mu = 10^{-4}$ per locus per generation. Alleles at 20/20/460 loci were beneficial/deleterious/neutral with selection coefficients +0.025/-0.025/0 respectively. The fitness landscape had a repeating comb-like structure shown in Supplementary **Fig. S1B**. One founder sequence was used to generate each population, which was allowed to evolve for 400 generations and sampled at $\Delta t = 50$ generations. Results are for MPL-R with $M = 3$ bootstrap samples each of size $R_b = R/3$ reads (coverage 1500X), where $R$ is the total number of reads available at each sampled time-point. The error bars indicate the standard error of the mean.

| Mut nt / Ref nt | A | C | G | T | - (gap) |
|---|---|---|---|---|---|
| A | 0 | $9.0 \times 10^{-7}$ | $6.0 \times 10^{-6}$ | $7.0 \times 10^{-7}$ | $1.0 \times 10^{-9}$ |
| C | $5.0 \times 10^{-6}$ | 0 | $5.0 \times 10^{-7}$ | $1.2 \times 10^{-5}$ | $1.0 \times 10^{-9}$ |
| G | $1.6 \times 10^{-5}$ | $1.0 \times 10^{-7}$ | 0 | $2.0 \times 10^{-6}$ | $1.0 \times 10^{-9}$ |
| T | $3.0 \times 10^{-6}$ | $1.0 \times 10^{-5}$ | $3.0 \times 10^{-6}$ | 0 | $1.0 \times 10^{-9}$ |
| - (gap) | $1.0 \times 10^{-9}$ | $1.0 \times 10^{-9}$ | $1.0 \times 10^{-9}$ | $1.0 \times 10^{-9}$ | 0 |

**Table S1. Mutation rates used in the analysis of HIV-1 patient data.** Each entry represents the mutation rate from the nucleotide in the row to the nucleotide in the column.

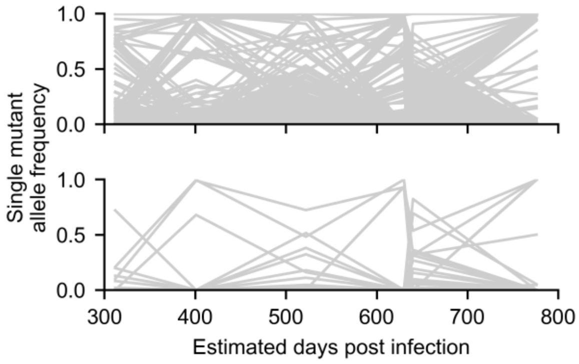


**Fig S3. SARS-CoV-2 data indicates presence of linkage.** Single mutant allele frequency trajectories obtained from the complete SARS-CoV-2 genome (*top* panel) and RBD (*bottom* panel) show multiple cooccurring mutations, suggesting the presence of genetic linkage.

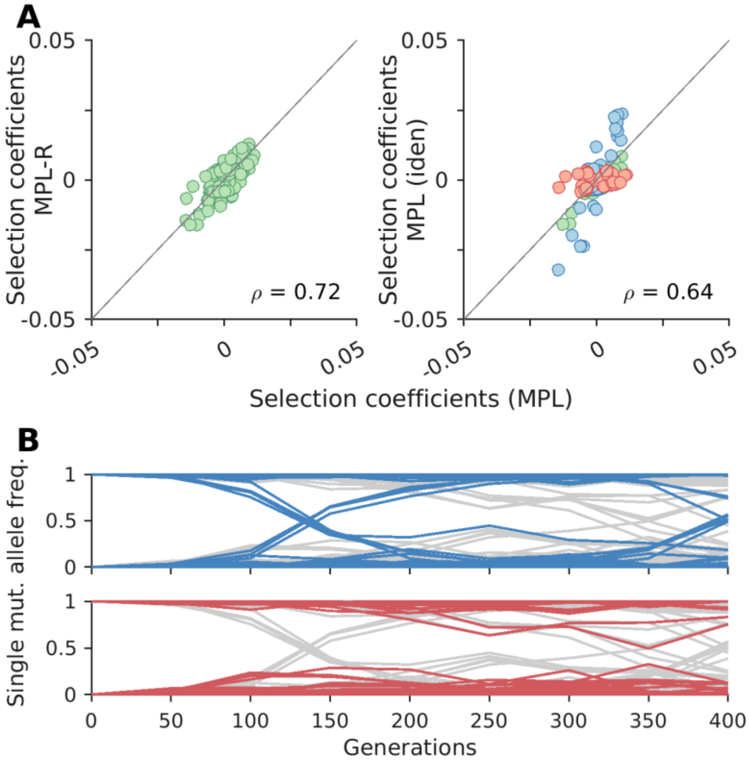


**Fig. S4. MPL (identity covariance) gets outcompeted by MPL-R for loci evolving under genetic linkage: (A)** Selection coefficients estimated from MPL-R and MPL (*left* panel) showed higher correlation (Pearson correlation coefficient $\rho = 0.72$ with p-value = $1.49 \times 10^{-58}$) compared to the selection coefficients obtained from MPL (identity covariance) and MPL (*right* panel) (Pearson correlation coefficient $\rho = 0.64$ with p-value = $1.28 \times 10^{-43}$). MPL (identity covariance) overestimates some selection coefficients (blue) and underestimates some (red). **(B)** Single mutant allele frequency trajectories for overestimated selection coefficients show a rising/falling trend (*top* panel). Single mutant allele frequency trajectories of underestimated selection coefficients start and end near a boundary. Results shown here are for a typical Monte Carlo run which consisted of evolving populations of $N = 1000$ individuals of $L = 500$ bi-allelic (WT and mutant) loci, with equal forward and backward mutation probabilities set to $\mu = 10^{-4}$ per locus per generation. Alleles at 20/20/460 loci were beneficial/deleterious/neutral with selection coefficients +0.025/-0.025/0 respectively. The fitness landscape had a repeating comb-like structure shown in Supplementary **Fig. S1B**. One founder sequence was used to generate each population, which was allowed to evolve for 400 generations and sampled at $\Delta t = 50$ generations. Results for MPL-R are with $M = 3$ bootstrap samples each of size $R_b = R/3$ reads (coverage 1500X), where $R$ is the total number of reads available at each sampled time-point.

### SI. 1 Comparison between MPL-R and MPL (identity covariance)

The selection coefficients estimated by MPL-R are relatively strongly correlated with those estimated by MPL (Pearson correlation coefficient $\rho = 0.72$ with p-value $= 1.49 \times 10^{-58}$) (Supplementary **Fig. S4A** (*left* panel)) as compared to the selection coefficients estimated by MPL (identity covariance) (Supplementary **Fig. S4A** (*right* panel)) (Pearson correlation coefficient $\rho = 0.64$ with p-value $= 1.28 \times 10^{-43}$). We can explain the performance difference between MPL-R and MPL (identity covariance) by analyzing the loci for which the latter performs comparatively worse. The relative error between the estimates of MPL (identity covariance) and MPL, and between MPL-R and MPL is given by

$$e_{i,\mathrm{MPL\ (iden)}} = |(s_{i,\mathrm{MPL}} - s_{i,\mathrm{MPL\ (iden)}})/s_{i,\mathrm{MPL}}|,$$

and

$$e_{i,\mathrm{MPL-R}} = |(s_{i,\mathrm{MPL}} - s_{i,\mathrm{MPL-R}})/s_{i,\mathrm{MPL}}|$$

respectively. $s_{i,\mathrm{MPL\ (iden)}}$, $s_{i,\mathrm{MPL-R}}$ , and $s_{i,\mathrm{MPL}}$ denote the selection coefficient for the mutant allele at locus $i$ estimated by MPL (identity covariance), MPL-R, and MPL respectively, $e_{i,\mathrm{MPL\ (iden)}}$ and $e_{i,\mathrm{MPL-R}}$ represent the relative error in the estimate of MPL (identity covariance) and MPL-R with respect to the estimate of MPL at locus $i$. $|\cdot|$ represents the absolute value. We first isolate all loci for which $e_{i,\mathrm{MPL\ (iden)}} > e_{i,\mathrm{MPL-R}}$, and then further classify those loci based on whether $s_{i,\mathrm{MPL\ (iden)}}$ is over or underestimated relative to $s_{i,\mathrm{MPL}}$. The scatterplot in Supplementary **Fig. S4A** (*right* panel) shows examples of overestimation $(|s_{i,\mathrm{MPL\ (iden)}}| > |s_{i,\mathrm{MPL}}|)$ highlighted as blue and examples of underestimation $(|s_{i,\mathrm{MPL\ (iden)}}| < |s_{i,\mathrm{MPL}}|)$ highlighted as red. The corresponding trajectories for the over and underestimated selection coefficients are presented in the same color in Supplementary **Fig. S4B** against all other trajectories colored gray. It can be observed that MPL (identity covariance) mostly overestimates the selection coefficients of loci for which the single mutant allele frequency trajectories are increasing or decreasing with time (Supplementary **Fig. S4B** (*top* panel)). This corresponds to the situation in which only looking at the term $x_j(t_K) - x_j(t_0)$ in **Equation 1** suggests a large value of the selection coefficient because of the large difference between the initial and final values of the trajectory, but the actual value of the selection coefficient, after correction for linkage, is small. These trajectories represent single mutant alleles that are inherently neutral but are hitchhiking on neighboring alleles because of genetic linkage. As MPL (identity covariance) does not correct for linkage, it overestimates the selection coefficients at these loci. It can also be observed that MPL (identity covariance) mostly underestimates the selection coefficients of loci for which the single mutant allele frequency trajectories start near the fixation/elimination boundary, increase or decrease for some time, and then end up at or close to the initial fixation/elimination boundary again (Supplementary **Fig. S4B** (*bottom* panel)). This corresponds to the situation in which only looking at the term $x_j(t_K) - x_j(t_0)$ in **Equation 1** suggests the selection coefficient to be zero because the initial and final values of the trajectories are similar, but the actual value of the selection coefficient, after correction for linkage, is non-zero. These trajectories represent single mutant alleles that are under positive selection but get outcompeted by alleles with stronger values of the selection coefficient (clonal interference) or are under negative selection but rise because of genetic linkage with positively selected neighboring alleles. Again, as MPL (identity covariance) does not correct for linkage, it underestimates the selection coefficients at these loci.

Trajectories driven by stronger selection coefficients can be hypothesized to increase the difference between MPL-R and MPL (identity covariance). To demonstrate the effect of the selection strength on the estimates of MPL-R and MPL (identity covariance), we compare the results of the datasets with $s = 0.075$ and $s = 0.025$ respectively. We can observe that the difference between MPL-R and MPL (identity covariance) is greater when $s = 0.075$ as compared to when $s = 0.025$ (Supplementary **Fig. S5**). The dataset with $s = 0.075$ has a higher magnitude of the selection coefficients, and when MPL (identity covariance) underestimates a large selection coefficient, the error is of a greater magnitude compared to the error that occurs when underestimates the selection coefficients for the dataset with $s = 0.025$.

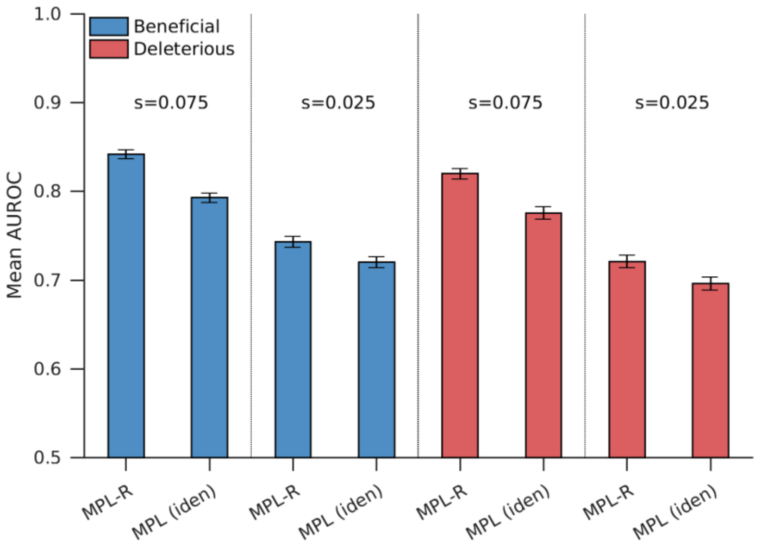


**Fig. S5. Performance improvement of MPL-R over MPL (identity covariance) is most significant for stronger selection coefficients:** Plot of the mean AUROC for beneficial loci (*left* sections) and deleterious loci (*right* sections) shows that the difference between the classification accuracy of MPL-R and MPL (identity covariance) increase with increase in the level of linkage. Results are shown for 100 Monte Carlo runs. Each Monte Carlo run consisted of evolving populations of $N = 1000$ individuals of $L = 500$ bi-allelic (WT and mutant) loci, with equal forward and backward mutation probabilities set to $\mu = 10^{-4}$ per locus per generation. Alleles at 20/20/460 loci were beneficial/deleterious/neutral with selection coefficients +0.025/-0.025/0 or +0.075/-0.075/0 respectively. The fitness landscape had a repeating comb-like structure shown in Supplementary **Fig. S1B**. One founder sequence was used to generate each population, which was allowed to evolve for 400 generations and sampled at $\Delta t = 50$ generations. Results for MPL-R are with $M = 3$ bootstrap samples each of size $R_b = R/3$ reads (coverage 1500X), where $R$ is the total number of reads available at each sampled time-point. The error bars indicate the standard error of the mean.

### SI. 2 Naïve estimation of ICM from reads

Here we describe a counterintuitive observation we made about the behavior of the ICM used in MPL (banded). MPL (banded) incorporates more information about the covariance between mutant allele frequencies than MPL (identity covariance), leading to a better estimate of the ICM. This can be seen from Supplementary **Fig. S6A**, which plots the normalized root mean square error (NRMSE) of the banded ICM as a function of increasing read-length (i.e., band length). Here, we calculate the NRMSE as

$$NRMSE = ||ICM_{\text{full}} - ICM_{\text{band}}||/||ICM_{\text{full}}||,$$

where $ICM_{\text{full}}$ denotes the ICM computed from full-length sequences, $ICM_{\text{band}}$ denotes the ICM of the banded case computed from the reads, and $||\cdot||$ denotes the Frobenius norm. When the read-length is zero in Supplementary **Fig. S6A**, the ICM of the banded case (solid line) reduces to the diagonal matrix of the identity covariance case (dashed line). While when the read-length is equal to the sequence length ($L = 500$) in Supplementary **Fig. S6A**, $ICM_{\text{band}} = ICM_{\text{full}}$. That is, incorporating more data (due to increased read length) successively improves the estimate of the ICM.

However, note that the estimate of the selection coefficients in **Equation 1** of the manuscript involves an inverse of the regularized ICM. Plotting the NRMSE of the inverse of the regularized ICM as a function of read length shows that the minimum NRMSE occurs for the $ICM_{\text{band}} = ICM_{\text{full}}$ case as expected (Supplementary **Fig. S6B**). Interestingly, we see that the NRMSE of the identity covariance case is lower than the banded covariance case. This is likely due to the non-linear nature of the inverse function. This inaccuracy in the inverse of the regularized ICM in the banded case at intermediate read-lengths leads to poor performance of MPL (banded) in classifying beneficial/deleterious mutants from the rest (Supplementary **Fig. S6C** and **D**). While it would be interesting to study this phenomenon in more detail, a more rigorous analysis is beyond the scope of the current manuscript.

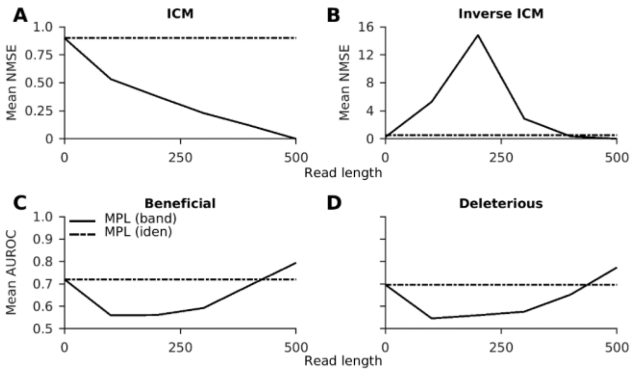


**Fig. S6. MPL (banded) performs poorly regardless of read length: (A)** The mean NRMSE of the ICM decreases with an increase in the read length. **(B)** The mean NRMSE of the inverse ICM first increases, then decreases with an increase in the read length. Plots of the mean AUROC for beneficial mutant alleles **(C)** and deleterious mutant alleles **(D)** show that MPL (banded) has poor classification performance compared to MPL (identity covariance). Results are shown for 100 Monte Carlo runs. Each Monte Carlo run consisted of evolving populations of $N = 1000$ individuals of $L = 500$ bi-allelic (WT and mutant) loci, with equal forward and backward mutation probabilities set to $\mu = 10^{-4}$ per locus per generation. Alleles at $20/20/460$ loci were beneficial/deleterious/neutral with selection coefficients $+0.025/-0.025/0$ respectively. The fitness landscape had a repeating comb-like structure shown in Supplementary **Fig. S1B**. One founder sequence was used to generate each population, which was allowed to evolve for 400 generations and sampled at $\Delta t = 50$ generations.

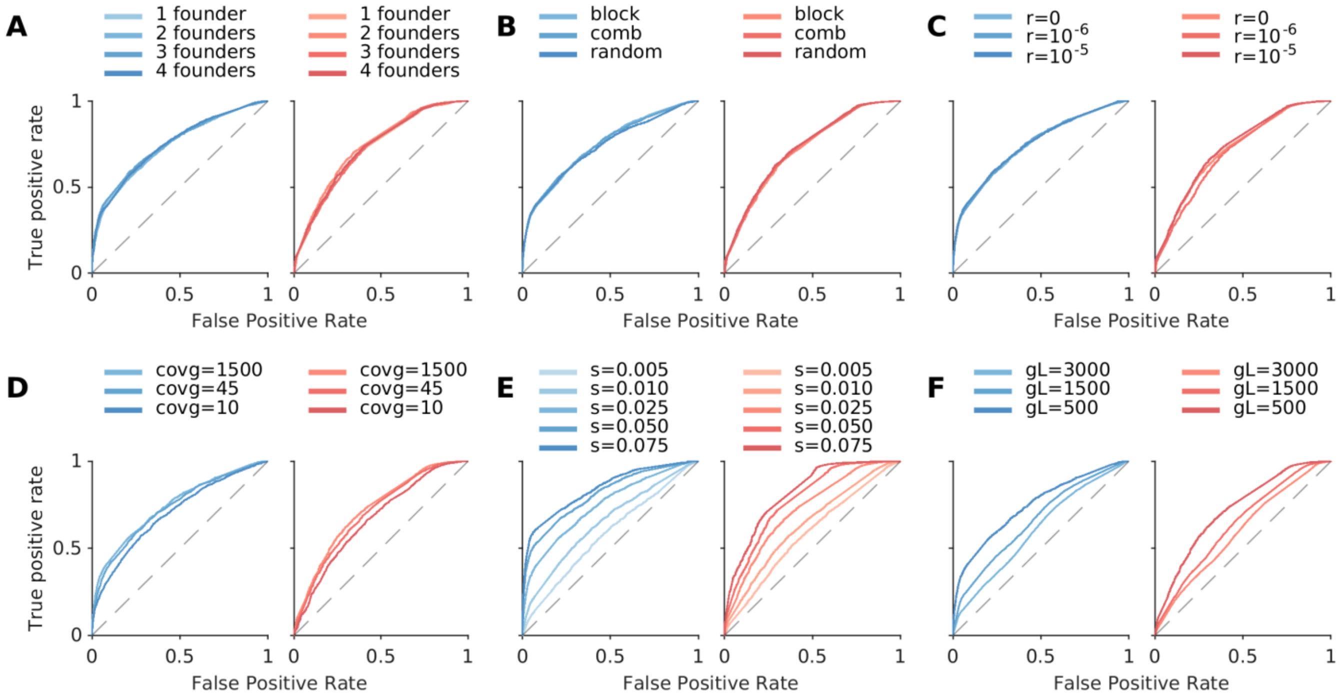


**Fig. S7. MPL-R consistently performs better than a random predictor when applied to datasets with changing diversity in data, distribution of fitness effect across the length of the sequence, recombination probability, strength of selection, length of the genome, and coverage.** The classification performance of MPL-R quantified as the mean receiver operating characteristic curve (ROC) consistently remained better than a random predictor for **(A)** increase in population diversity, controlled here by varying the number of founder strains (frequency of each founder ≥ 10%), **(B)** various structures of the underlying fitness landscape, **(C)** increase in recombination probability, **(D)** sequencing coverage, **(E)** decrease in selection strength, and **(F)** increase in the length of the genome. The structure of the data sets "comb", "block", and "random" is presented in Supplementary **Fig. S1**. Results are shown for 100 Monte Carlo runs. Each Monte Carlo run consisted of evolving populations of $N = 1000$ individuals of $L = 500$ bi-allelic (WT and mutant) loci, with equal forward and backward mutation probabilities set to $\mu = 10^{-4}$ per locus per generation. Unless mentioned otherwise, alleles at $20/20/460$ loci were beneficial/deleterious/neutral with selection coefficients $+0.025/-0.025/0$ respectively. Unless mentioned otherwise, the fitness landscape had a repeating comb-like structure shown in Supplementary **Fig. S1B**. Unless mentioned otherwise, one founder sequence was used to generate each population, which was allowed to evolve for 400 generations and sampled at $\Delta t = 50$ generations. Results are for MPL-R with $M = 3$ bootstrap samples each of size $R_b = R/3$ reads (coverage 1500X), where $R$ is the total number of reads available at each sampled time-point.

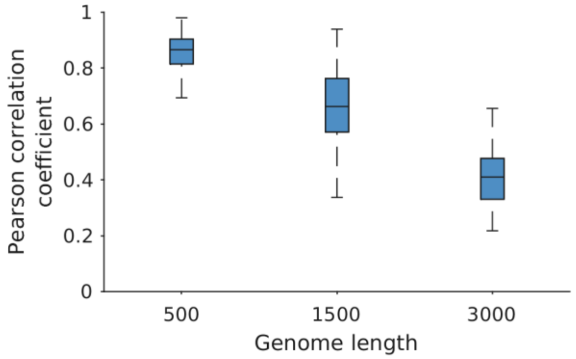


**Fig. S8. Increasing the genome length decreases the accuracy of the inferred covariance values.** The Pearson correlation coefficient between the reconstructed and ground truth covariance values indicate a consistent decreasing trend with the increase in genome length. The results are plotted for 100 Monte Carlo runs, each Monte Carlo run consisted of evolving populations of $N=$ 1000 individuals of bi-allelic (WT and mutant) loci of the indicated genome length, with equal forward and backward mutation probabilities set to $\mu = 10^{-4}$ per locus per generation. Alleles at 20/20/460, 60/60/1380, and 120/120/1260, loci were beneficial/deleterious/neutral with selection coefficients +0.025/-0.025/0 respectively. The fitness landscape had a repeating comb-like structure shown in Supplementary **Fig. S1B**. One founder sequence was used to generate each population, which was allowed to evolve for 400 generations and sampled at $\Delta t = 50$ generations. Results are for MPL-R with $M = 3$ bootstrap samples each of size $R_b = R/3$ reads (coverage 1500X), where $R$ is the total number of reads available at each sampled time-point.

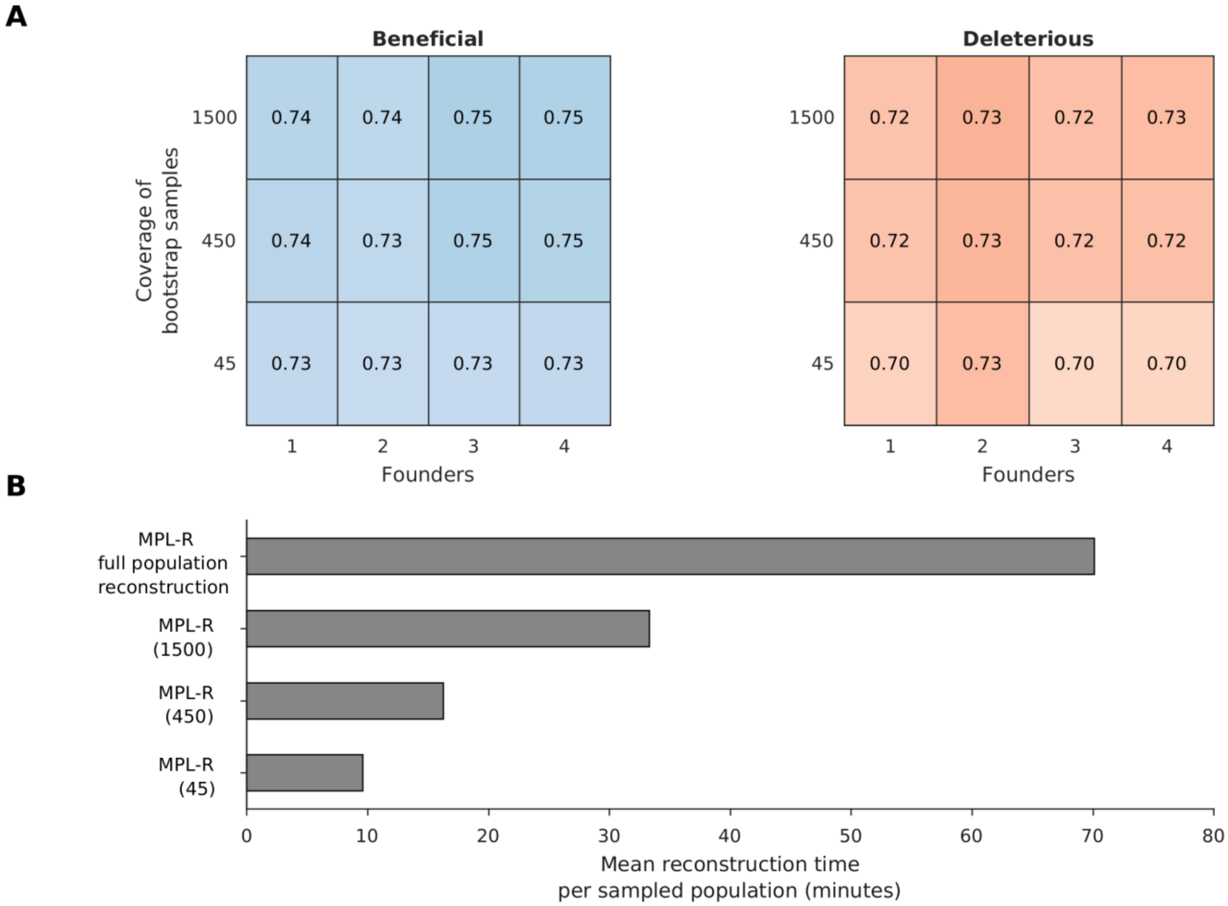


**Fig. S9. Decrease in the size of the bootstrap samples decreases the wall clock time with a gradual decrease in the classification accuracy. (A)** Comparison of the mean AUROC of MPL-R for different sizes of the bootstrap samples. The bootstrap sample sizes are indicated in terms of the average coverage. The classification accuracy is robust to the bootstrap sample size. **(B)** Average reconstruction time of the population at each time point for MPL-R without bagging, and MPL-R with different sizes of the bootstrap samples indicated in the parenthesis. Results of MPL-R are for $M = 3$ bootstrap samples. Results are shown for 100 Monte Carlo runs. Each Monte Carlo run consisted of evolving populations of $N = 1000$ individuals of $L = 500$ bi-allelic (WT and mutant) loci, with equal forward and backward mutation probabilities set to $\mu = 10^{-4}$ per locus per generation. Alleles at 20/20/460 loci were beneficial/deleterious/neutral with selection coefficients +0.025/-0.025/0 respectively. The fitness landscape had a repeating comb-like structure shown in Supplementary **Fig. S1B**. One founder sequence was used to generate each population, which was allowed to evolve for 400 generations and sampled at $\Delta t = 50$ generations.

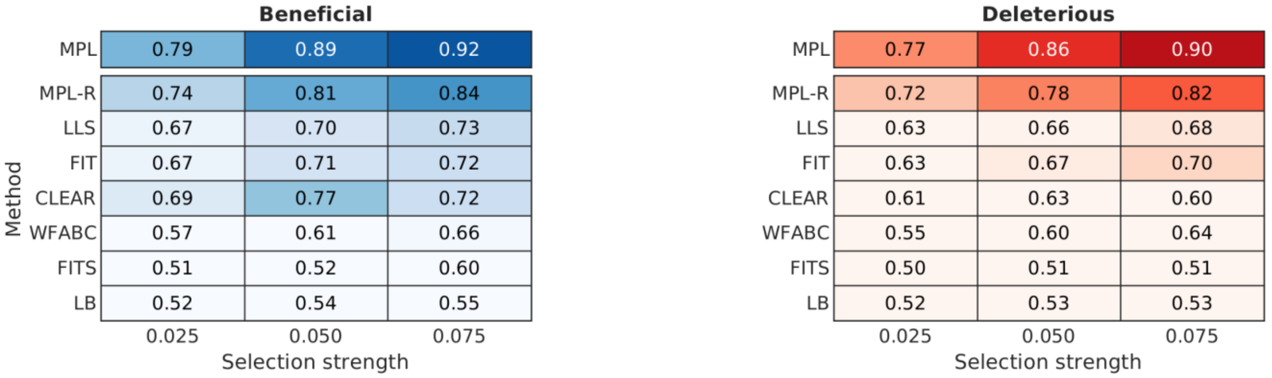


**Fig. S10. MPL-R consistently performs better than available inference methods.** Comparison of classification performance (measured in AUROC) of MPL-R and state-of-the-art methods for varying selection strength. A method with complete knowledge of all double mutant allele frequencies (MPL) is presented as a benchmark. Results for MPL-R are with $M = 3$ bootstrap samples each of size $R_b = R/3$ reads (coverage 1500X), where $R$ is the total number of reads available at each sampled time-point Results on the rest of the methods are on the single mutant allele frequencies computed from the short reads obtained from the ART simulator. Results are shown for 100 Monte Carlo runs. Each Monte Carlo run consisted of evolving populations of $N = 1000$ individuals of $L = 500$ bi-allelic (WT and mutant) loci, with equal forward and backward mutation probabilities set to $\mu = 10^{-4}$ per locus per generation. Alleles at 20/20/460 loci were beneficial/deleterious/neutral with selection coefficients +0.025/-0.025/0 respectively. The fitness landscape had a repeating comb-like structure shown in Supplementary **Fig. S1B**. One founder sequence was used to generate each population, which was allowed to evolve for 400 generations and sampled at $\Delta t = 50$ generations.

### SI. 3 Comparison with existing time-series methods

The existing time-series methods infer selection based on the rate of change (derivative) of single mutant allele frequencies. Such a model assumes selection as the driving force behind changes in the single mutant allele frequency, and selection inference boils down to the inverse problem of observing changes in the single mutant allele frequencies to quantify the magnitude and sign of selection. This approach will infer the selection coefficients as positive when the single mutant allele frequencies of the mutant alleles undergo an increase as evolution progresses, and negative if the single mutant allele frequencies decrease with time. This methodology does not explicitly model changes in the single mutant allele frequencies due to mutations, and may not be well-suited to analyze data that incorporates mutation. In all our tests, we simulated random mutations in the population, which may explain the degradation in performance of methods based on the rate of change of single mutant allele frequencies. In the following, we shall briefly describe the working principle of each of the competing methods and present an empirical test showing how the incorporation of mutation in the estimation yields better performance.

The method LLS (Taus *et al.*, 2017) logit-transforms the single mutant allele frequency trajectories and fits a linear model with least squares regression (LLS) to the transformed trajectories. The slope of the fit gives the selection coefficient. CLEAR (Iranmehr *et al.*, 2017) and WFABC (Foll *et al.*, 2015) both model the change in the single mutant allele frequencies as a function of the selection coefficients (s) and the dominance parameter (h). For CLEAR, the calculation of the selection coefficients is performed via maximum likelihood estimation given the observed single mutant allele frequencies. WFABC infers the selection coefficients via approximate Bayesian computation (ABC) (Sunnåker *et al.*, 2013) in which repeated simulations are performed to approximate the likelihood function. FITS (Zinger *et al.*, 2019) is also based on Approximate Bayesian Computation (ABC). The method FIT (Feder *et al.*, 2014) infers the selection coefficients based on the increments in the single mutant allele frequency trajectories at different time intervals. The assumption is that if the single mutant allele frequency increments over time belong to a zero-mean normal distribution, the underlying selection is neutral, as there is no net increase/decrease in the single mutant allele frequency trajectory.

In summary, all these methods have different working principles, implementations, and parameters optimized for the implementations. It is difficult to make an exact comparison about what factors contribute to the difference in performance with MPL (identity covariance), but there is one fundamental difference between the working principle of MPL (identity covariance) and the rest of the methods. MPL (identity covariance) not only models the change in the single mutant allele frequency trajectories as a function of selection, but it also considers the change in single mutant allele frequency trajectories because of mutation via the mutational flux term (**Equation 1**). Intuitively, the mutational flux of a mutation quantifies the tendency of other alleles to acquire that mutation. For a beneficial mutant allele, selective pressure favors the mutation of WT alleles into the mutant, hence the mutational flux is positive. The mutation flux for neutral and deleterious mutant

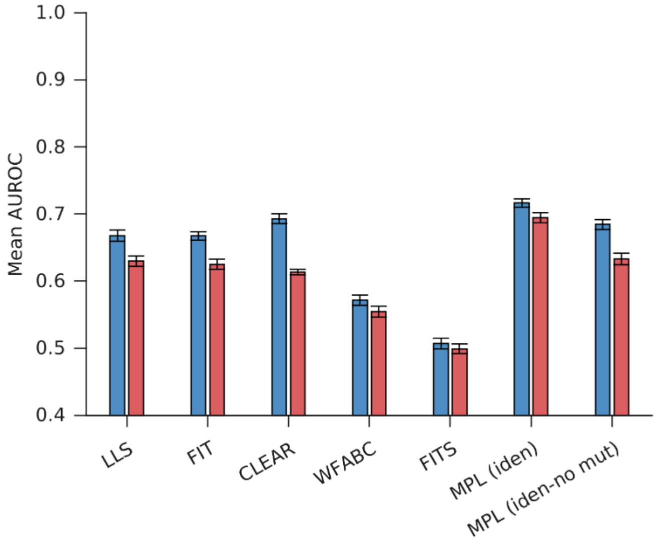


**Fig. S11. The performance of MPL (identity covariance) degrades when mutational flux is ignored:** Plot of the mean AUROC for beneficial loci (*blue*) and deleterious loci (*red*) shows that the performance of MPL (identity covariance-no mutational flux) degrades because of ignoring mutational flux. Results are shown for 100 Monte Carlo runs. Each Monte Carlo run consisted of evolving populations of $N = 1000$ individuals of $L = 500$ bi-allelic (WT and mutant) loci, with equal forward and backward mutation probabilities set to $\mu = 10^{-4}$ per locus per generation. Alleles at 20/20/460 loci were beneficial/deleterious/neutral with selection coefficients +0.025/-0.025/0 respectively. The fitness landscape had a repeating comb-like structure shown in Supplementary **Fig. S1B**. One founder sequence was used to generate each population, which was allowed to evolve for 400 generations and sampled at $\Delta t = 50$ generations. The error bars indicate the standard error of the mean.

alleles is zero and negative respectively based on a similar argument. So, even in the scenarios where the change in the single mutant allele frequency trajectories is small, the mutational flux still captures the effect of selection. To demonstrate the effect of mutational flux on the classification performance, we compare the results of MPL (identity covariance) with another implementation in which the mutational flux is not considered, hereby referred to as MPL (identity covariance-no mutational flux). Because only the change in single mutant allele frequencies is used to infer selection coefficients for MPL (identity covariance-no mutational flux), the method makes more mistakes compared to MPL (identity covariance), which takes into account the mutational flux. This resulted in degraded performance of MPL (identity covariance-no mutational flux), which was more comparable to that of CLEAR, which also doesn't take into account mutational flux (Supplementary **Fig. S11**). It is also important to mention here that incorporation of the effects of mutation into the estimator is one reason, but it may not be the only reason for the difference in performance in the methods. It is hard to make a direct comparison between these methods and pinpoint the exact reasons behind the differences in performance, but we have empirically shown that incorporation of mutation does seem to add statistical power to the estimation.

| | CD8+ T cell escape mutations | | | Nonsynonymous reversions outside CD8+ T cell epitopes | | | Nonsynonymous reversions within CD8+ T cell epitopes | | |
|---|---|---|---|---|---|---|---|---|---|
| | % | Fold enrichment | p-value | % | Fold enrichment | p-value | % | Fold enrichment | p-value |
| MPL-R | 40.5 | 23 | $< 10^{-13}$ | 13.5 | 21 | $< 10^{-5}$ | 10.8 | 358 | $< 10^{-8}$ |
| MPL | 34.3 | 18 | $< 10^{-9}$ | 17.1 | 27 | $< 10^{-7}$ | 11.4 | 382 | $< 10^{-8}$ |
| MPL (iden) | 28.6 | 14 | $< 10^{-7}$ | 22.9 | 33 | $< 10^{-9}$ | 11.4 | 382 | $< 10^{-8}$ |

**Table S2. MPL-R reports similar insights as MPL.** Percentage of mutations, fold enrichment, and p-values reported by MPL-R, MPL, and MPL (identity covariance) are given for mutations in the listed categories in the top 5% mutations with the strongest selection coefficients.

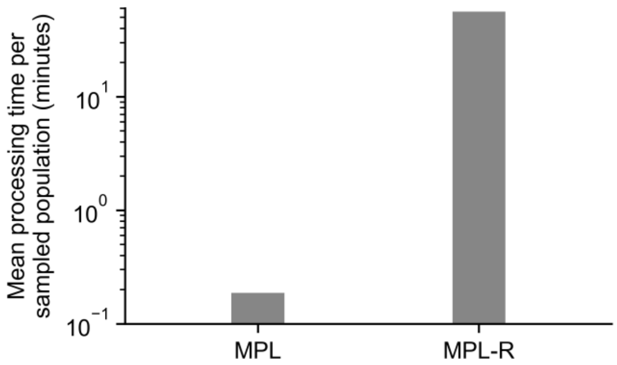


**Fig. S12. MPL-R completes the analysis on the HIV-1 data in a reasonable amount of time.** The time required by MPL-R to analyze the HIV-1 data is less than an hour. Though it takes much longer than MPL, it needs to be taken into consideration that full-sequence data required by MPL may not always accessible to the end user, and MPL-R might the only available tool that can perform inference.

| Mutation | Experimentally reported phenotypic effects |
|---|---|
| S371F | Antibody escape (Cao *et al.*, 2022a, 2022b) |
| F377L | Antibody escape (Huo *et al.*, 2021) |
| D405N | Antibody escape (Cao *et al.*, 2022b) |
| K417N | Antibody escape (Yuan *et al.*, 2021, Wang *et al.*, 2021) |
| N440K | Antibody escape (Rani *et al.*, 2021) |
| N450K | Antibody escape (Liu *et al.*, 2021) |
| Q493R | Antibody escape (Focosi *et al.*, 2021) |
| Q498R | Antibody escape (Cui *et al.*, 2022), host-binding enhancement (Hong *et al.*, 2022) |
| Y505H | Antibody escape (Wang *et al.*, 2023) |
| G339D | T-cell response weakening (Li *et al.*, 2022), antibody escape (Cao *et al.*, 2022a) |
| N354K | Immunogenicity reduction and antibody escape (Liu *et al.*, 2024) |
| L452Q | Stability enhancement (Starr *et al.*, 2022), antibody escape (Li *et al.*, 2020, Cao *et al.*, 2022b), infectivity enhancement (Deng *et al.*, 2021) |
| V367F | Infectivity enhancement (Ou *et al.*, 2021) |
| S375F | Infectivity enhancement (Kimura *et al.*, 2022) |
| S373P | Host-binding enhancement (Zheng *et al.*, 2023) |
| N501Y | Host-binding enhancement (Tian, 2021) |
| T376A | Spike cleavage reduction (Hu *et al.*, 2022) |

**Table S3. Mutations inferred as beneficial by MPL-R map to known beneficial mutations.** Nonsynonymous mutations in the RBD which are inferred as beneficial by MPL-R and neutral or deleterious by MPL (identity covariance) map to mutations reported to be beneficial to the virus.

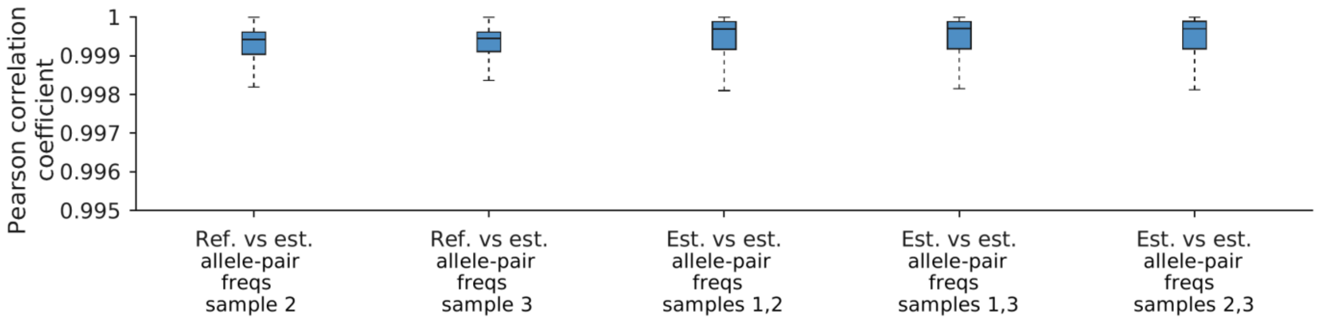


**Fig. S13. The reconstructed double mutant allele frequencies are accurate.** The double mutant allele frequencies calculated from the reconstructed and actual populations are highly correlated. The double mutant allele frequencies calculated from different bootstrap samples also exhibit strong correlation. Results are for MPL-R with $M = 3$ bootstrap samples each of size $R_b = R/3$ reads (coverage 1500X), where $R$ is the total number of reads available at each sampled time-point. Results are shown for 100 Monte Carlo runs. Each Monte Carlo run consisted of evolving populations of $N = 1000$ individuals of $L = 500$ bi-allelic (WT and mutant) loci, with equal forward and backward mutation probabilities set to $\mu = 10^{-4}$ per locus per generation. Alleles at 20/20/460 loci were beneficial/deleterious/neutral with selection coefficients +0.025/-0.025/0 respectively. The fitness landscape had a repeating comb-like structure shown in Supplementary **Fig. S1B**. One founder sequence was used to generate each population, which was allowed to evolve for 400 generations and sampled at $\Delta t = 50$ generations.

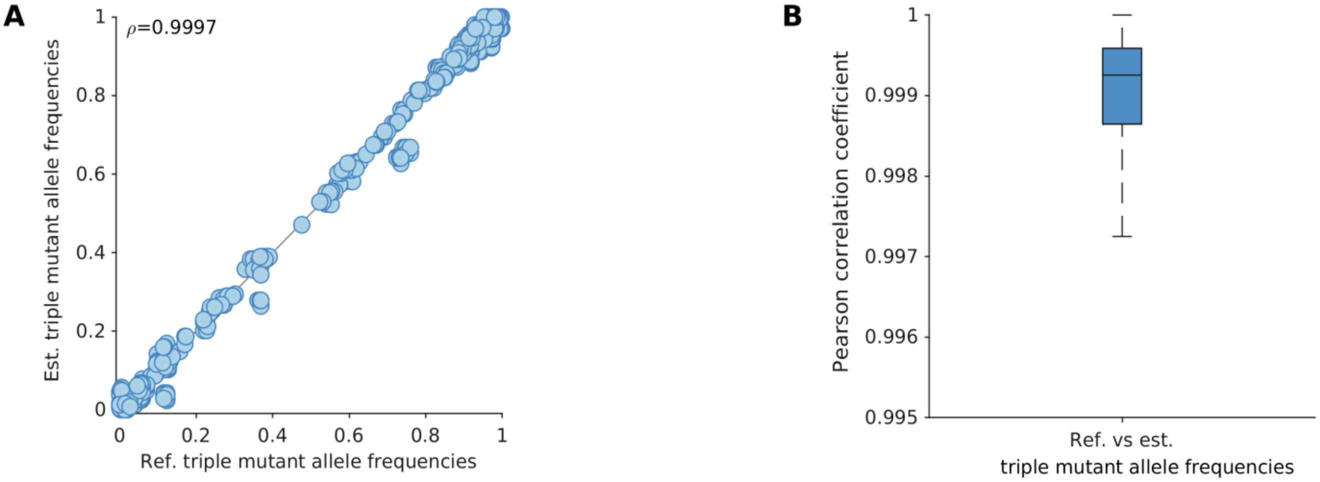


**Fig. S14. MPL-R faithfully reconstructs the triple mutant allele frequencies. (A)** Triple mutant allele frequencies calculated from the reconstructed population are highly correlated with the actual triple mutant allele frequencies (Pearson correlation coefficient $\rho = 0.9997$ with p-value $< 10^{-100}$). **(B)** The summary statistics of the Pearson correlation coefficient between the estimated and actual triple mutant allele frequencies indicate a consistent trend across all Monte Carlo runs. Results are for $M = 3$ bootstrap samples each of size $R_b = R/3$ reads (coverage 1500X), where $R$ is the total number of reads available at each sampled time-point. Results are shown for 100 Monte Carlo runs. Each Monte Carlo run consisted of evolving populations of $N = 1000$ individuals of $L = 500$ bi-allelic (WT and mutant) loci, with equal forward and backward mutation probabilities set to $\mu = 10^{-4}$ per locus per generation. Alleles at 20/20/460 loci were beneficial/deleterious/neutral with selection coefficients +0.025/-0.025/0 respectively. The fitness landscape had a repeating comb-like structure shown in Supplementary **Fig. S1B**. One founder sequence was used to generate each population, which was allowed to evolve for 400 generations and sampled at $\Delta t = 50$ generations.

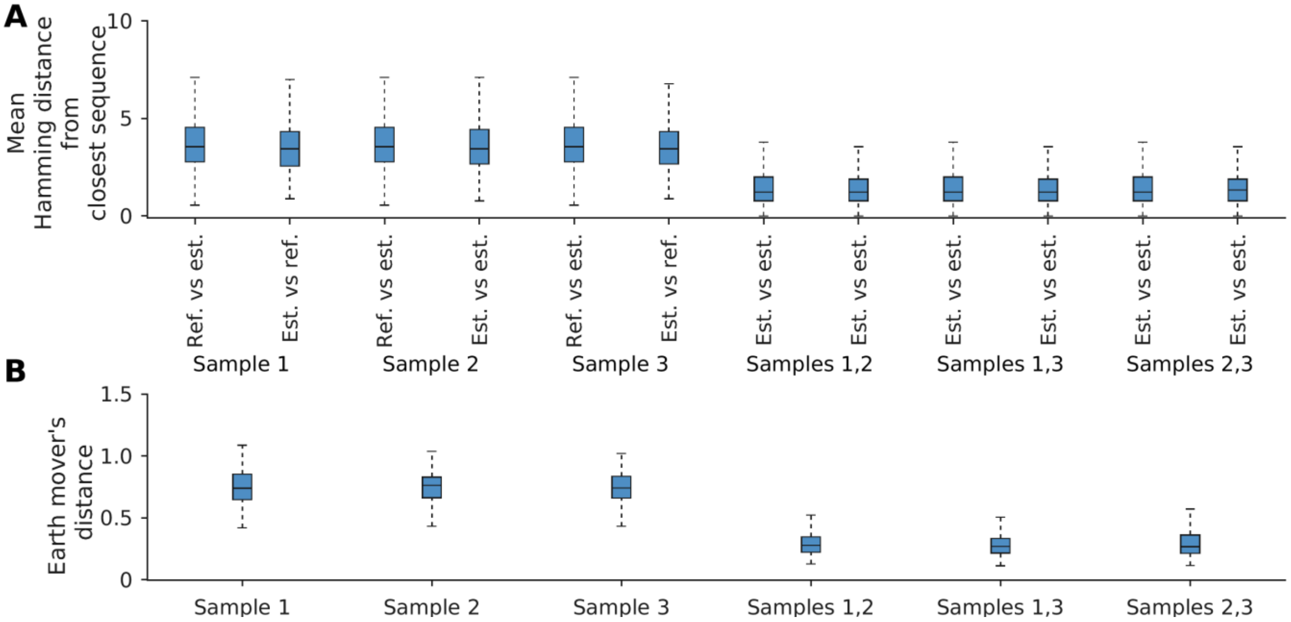


**Fig. S15. The reconstruction performed via Quasirecomb is accurate. (A)** The mean Hamming distance between the actual Wright Fisher population and the closest sequences in the reconstructed population and vice versa is small. Sequences in the reconstructed populations differ from each other by a small Hamming distance as well. **(B)** The earth mover's distance between the reference Wright Fisher population and reconstructed populations and between the reconstructed populations is small. Results are for $M = 3$ bootstrap samples each of size $R_b = R/3$ reads (coverage 1500X), where $R$ is the total number of reads available at each sampled time-point. Results are shown for 100 Monte Carlo runs. Each Monte Carlo run consisted of evolving populations of $N = 1000$ individuals of $L = 500$ bi-allelic (WT and mutant) loci, with equal forward and backward mutation probabilities set to $\mu = 10^{-4}$ per locus per generation. Alleles at 20/20/460 loci were beneficial/deleterious/neutral with selection coefficients +0.025/-0.025/0 respectively. The fitness landscape had a repeating comb-like structure shown in Supplementary **Fig. S1B**. One founder sequence was used to generate each population, which was allowed to evolve for 400 generations and sampled at $\Delta t = 50$ generations.

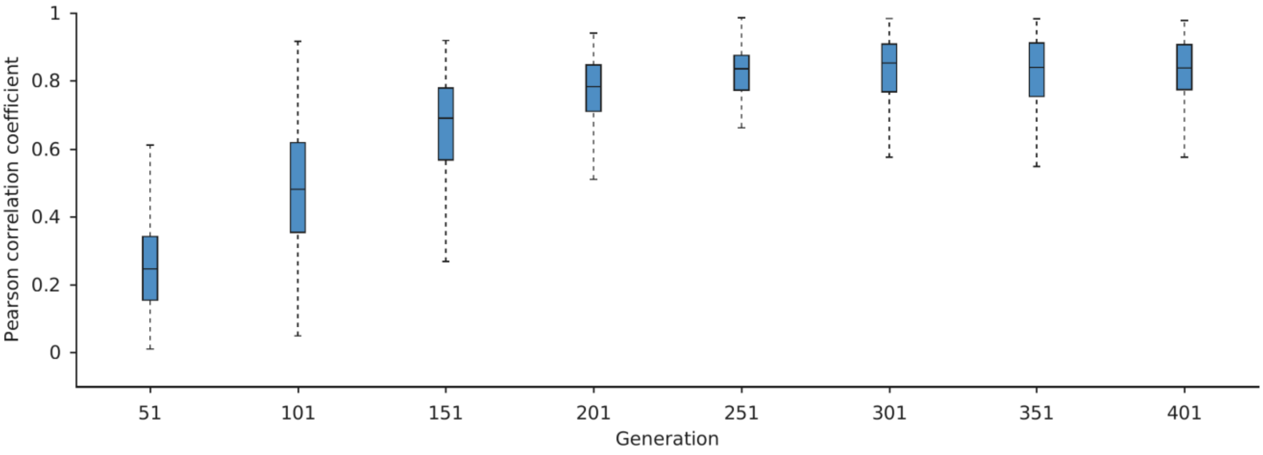


**Fig. S16. The covariance matrices of allele frequencies calculated using reconstructed double mutant allele frequencies at each time point are accurate.** The covariance matrices of allele frequencies calculated from the reconstructed and actual populations exhibit good correlation across the sampled time-points. The correlation at the initial time points is poor because the single mutant allele frequencies at these time points are small (**Fig. 2A**), and hence the covariance matrices have negligible magnitudes. Results are for $M = 3$ bootstrap samples each of size $R_b = R/3$ reads (coverage 1500X), where $R$ is the total number of reads available at each sampled time-point. Results are shown for 100 Monte Carlo runs. Each Monte Carlo run consisted of evolving populations of $N = 1000$ individuals of $L = 500$ bi-allelic (WT and mutant) loci, with equal forward and backward mutation probabilities set to $\mu = 10^{-4}$ per locus per generation. Alleles at 20/20/460 loci were beneficial/deleterious/neutral with selection coefficients +0.025/-0.025/0 respectively. The fitness landscape had a repeating comb-like structure shown in Supplementary **Fig. S1B**. One founder sequence was used to generate each population, which was allowed to evolve for 400 generations and sampled at $\Delta t = 50$ generations.

### References

Cao, Y. *et al.* (2022a) Omicron escapes the majority of existing SARS-CoV-2 neutralizing antibodies. *Nature*, **602**, 657-663.

Cao, Y. *et al.* (2022b) BA. 2.12. 1, BA. 4 and BA. 5 escape antibodies elicited by Omicron infection. *Nature*, **608**, 593-602.

Cui, Z. *et al.* (2022) Structural and functional characterizations of infectivity and immune evasion of SARS-CoV-2 Omicron. *Cell*, **185**, 860-871.

Deng, X. *et al.* (2021) Transmission, infectivity, and neutralization of a spike L452R SARS-CoV-2 variant. *Cell*, **184**, 3426-3437.

Focosi, D. *et al.* (2021) Emergence of SARS-COV-2 spike protein escape mutation Q493R after treatment for COVID-19. *Emerg. Infect. Dis.*, **27**, 2728.

Hong, Q. *et al.* (2022) Molecular basis of receptor binding and antibody neutralization of Omicron. *Nature*, **604**, 546-552.

Hu, B. *et al.* (2022) Spike mutations contributing to the altered entry preference of SARS-CoV-2 omicron BA. 1 and BA. 2. *Emerg. Microbes Infect.*, **11**, 2275-2287.

Huo, J. *et al.* (2021) A potent SARS-CoV-2 neutralising nanobody shows therapeutic efficacy in the Syrian golden hamster model of COVID-19. *Nat. Commun.*, **12**, 5469.

Kimura, I. *et al.* (2022) The SARS-CoV-2 spike S375F mutation characterizes the Omicron BA. 1 variant. *iScience*, **25**.

Li, Q. *et al.* (2020) The impact of mutations in SARS-CoV-2 spike on viral infectivity and antigenicity. *Cell*, **182**, 1284-1294.

Li, Y. *et al.* (2022) T-cell responses to SARS-CoV-2 Omicron spike epitopes with mutations after the third booster dose of an inactivated vaccine. *J. Med. Virol.*, **94**, 3998-4004.

Liu, P. *et al.* (2024) Spike N354 glycosylation augments SARS-CoV-2 fitness for human adaptation through structural plasticity. *Natl. Sci. Rev.*, **11**, nwae206.

Liu, Z. *et al.* (2021) Identification of SARS-CoV-2 spike mutations that attenuate monoclonal and serum antibody neutralization. *Cell Host Microbe*, **29**, 477-488.

Ou, J. *et al.* (2021) V367F mutation in SARS-CoV-2 spike RBD emerging during the early transmission phase enhances viral infectivity through increased human ACE2 receptor binding affinity. *J. Virol.*, **95**, 10-1128.

Rani, P. *et al.* (2021) Symptomatic reinfection of SARS-CoV-2 with spike protein variant N440K associated with immune escape. *J. Med. Virol.*, **93**, 4163.

Starr, T. N. *et al.* (2022) Shifting mutational constraints in the SARS-CoV-2 receptor-binding domain during viral evolution. *Science*, **377**, 420-424.

Tian, F. (2021) N501Y mutation of spike protein in SARS-CoV-2 strengthens its binding to receptor ACE2. *eLife*, **10**, e69091.

Wang, W. B. *et al.* (2023) Identification of key mutations responsible for the enhancement of receptor-binding affinity and immune escape of SARS-CoV-2 Omicron variant. *J. Mol. Graph.*, **124**, 108540.

Wang, Z. *et al.* (2021) mRNA vaccine-elicited antibodies to SARS-CoV-2 and circulating variants. *Nature*, **592**, 616-622.

Yuan, M. *et al.* (2021) Structural and functional ramifications of antigenic drift in recent SARS-CoV-2 variants. *Science*, **373**, 818-823.

Zheng, B. *et al.* (2023) S373P mutation stabilizes the receptor-binding domain of the spike protein in omicron and promotes binding. *JACS Au*, **3**, 1902-1910.
